# Disrupted mannosidase enzymes in C9orf72 ALS/FTD

**DOI:** 10.64898/2026.10.30.763135

**Authors:** BN Ondatje, LM Gittings, KR Preller, E Alsop, K Garcia-Mansfield, N Hansen, B Lovell, R Sharma, IS Piras, M Martinez, B Meechoovet, A Bonfitto, S Song, J Levy, M Huentelman, P Pirrotte, K Van Keuren-Jensen, R Sattler

## Abstract

Amyotrophic lateral sclerosis (ALS) is an age-related neurodegenerative disorder resulting from the loss of motor neurons in the brain and spinal cord. Neuroinflammation, induced by glial cells including microglia (MG) and astrocytes, is an acknowledged aspect of this neurodegeneration. Generating induced pluripotent stem cell (iPSC) neuroinflammatory models of ALS may help elucidate non-neuronal mechanisms of disease. Here, we analyze human postmortem tissue and an iPSC derived glia-crosstalk model identifying a reduction in Golgi enzymes MAN1A1 and MAN1A2 in addition to other glycan enzymes from ALS-associated microglia and astrocytes. Impaired Golgi and glycan metabolism is linked to inflammation and other related ALS disease mechanisms including impaired lysosomal proteostasis, altered extracellular matrix, disrupted secretion, and altered cell-cell signaling.

## Introduction

ALS is a progressive neurodegenerative disease characterized by the loss of upper motor neurons of the cortex and brain stem, and lower motor neurons of the spinal cord^1^. Beyond motor neuron loss, ALS pathology is accompanied by prominent neuroinflammatory responses involving both microglia and astrocytes^2,3^, highlighting the contribution of non-neuronal cells to disease pathogenesis. ALS also extends beyond the motor system as many patients develop cognitive and behavioral impairments associated with frontal and temporal cortical degeneration, and ALS shares substantial clinical, pathological, and genetic overlap with frontotemporal dementia (FTD)^4,5^. Despite this complex and multisystem disease biology, limited therapeutic options exist for ALS patients^6^. Currently available treatments, including Riluzole, Edaravone, and the *SOD1* targeting antisense oligonucleotide tofersen (Qalsody)^7–11^, address selected aspects of neuronal dysfunction or disease-associated protein pathology but provide benefit to only subsets of patients or modestly alter disease progression. Furthermore, numerous clinical trials targeting individual disease mechanisms that drive ALS progression have failed to demonstrate meaningful efficacy^12–15^. Together, these limitations emphasize the need to better define the complex biology of this disease, including interactions among glial populations, to identify new druggable targets and tractable biomarkers.

Although ALS is predominantly sporadic, several genetic variants exist, the most prevalent of which is a G_4_C_2_ hexanucleotide repeat expansion (HRE) in the first intron of the *C9orf72* gene^4,5^. Pathogenic repeat lengths range from 38 to over 1500 repeats, with both loss-of-function (LOF) and gain-of-function (GOF) mechanisms found in neuronal and glial cell types^5^. While this subtype of ALS accounts for 40%-50% of inherited ALS and about 5%-10% of sporadic ALS, it shares a common pathomechanism accounting for 97% of the ALS population - the nuclear depletion and cytoplasmic accumulation of TAR DNA-binding protein 43 (TDP-43) - making this an ideal model system for mechanistic investigations with the goal of identifying new therapeutic targets and biomarkers ^16^.

One approach to identifying novel therapeutic targets and biomarkers is to investigate mechanisms of disease associated with non-neuronal cell types of the central nervous system (CNS). Neuroinflammation is a well-documented phenotype associated with neurodegeneration, with inflammatory signals detected early in disease progression^17–20^. Many studies have investigated how the cells associated with inflammatory responses in the CNS, microglia and astrocytes, interact with neurons, noting detrimental effects on the health of neurons^21–27^. Limited studies have determined the effects of these glial cells on each other, and how their interactions may contribute to disease onset and progression^3,28–35^. Understanding this relatively unexplored frontier for neurodegeneration opens an opportunity for identification of novel mechanisms of disease and innovative biomarkers.

In this study, we investigated the role of microglia-astrocyte crosstalk in C9orf72 (C9) ALS/FTD. Exploratory analysis of previously published single-nuclei RNA sequencing (snRNA-seq) from C9-ALS, C9-ALS/FTD, and healthy control human postmortem frontal cortex^36^ identified a reduction in the Golgi enzyme MAN1A1 in C9orf72-microglia, which also showed concurrent alterations in pathways linked to synaptic organization and astrocyte differentiation. These findings implicated altered glycosylation as a potentially disease-relevant process in microglia. These findings were validated in an induced pluripotent stem cell (iPSC) model of microglia-astrocyte crosstalk from C9orf72 ALS/FTD patients, which identified a reduction of the redundant, and developmentally active protein, MAN1A2. Additionally, we observed increases in thrombospondin 1 (THBS1)-vascular endothelial growth factor A (VEGFA) signaling in iPSC microglia (MG), triggered by secreted cues of C9orf72 ALS/FTD iPSC cortical astrocytes (CA). Furthermore, this work identified dysregulation of neuroinflammatory-linked neurodegenerative molecules including apolipoprotein E (APOE), amyloid precursor protein (APP), and presenilin 2 (PSEN2), highlighting overlapping neuroinflammatory signatures between ALS/FTD and Alzheimer’s Disease (AD)^19,37,38^. Lastly, ingenuity pathway-interactor analysis (IPA) suggested aberrations in signaling pathways between MG and CA, including APOE and transforming growth factor beta 1 (TGFβ1) consistent with observed molecular changes in APOE and THBS1. This work provides grounds for exploring identified mechanisms of disease for therapeutic targeting or establishment of novel disease biomarkers, and highlights Golgi function and glycosyl metabolism as pivotal components to disease and potential upstream dysregulation, as it plays an integral role in signal transduction, extracellular matrix, protein folding and stability, and immune regulation^39^, and has previously been associated with ALS/FTD^40–46^.

## Results

### Human postmortem microglia reveal impaired glycan processing and inflammatory activation in C9orf72 ALS/FTD

To determine whether alterations in glial molecular functions are associated with human C9orf72 ALS/FTD pathology, we first analyzed published single-nucleus RNA (snRNA) sequencing datasets from postmortem frontal cortex^36^. Microglia sub-clustering from 29 patients from the Gittings et al. publication^36^ led to four subclusters of distinct transcriptional states. Clusters 0 and 3 are enriched in inflammatory, complement, and apoptotic markers (Figure 1A & Figure S1A). Cluster 1 also indicates activation of complement signaling via elevated complement 3 expression (Figure 1A & Figure S1A), while cluster 2 indicates elevated complement *C1QA*, *APOE*, *STAT3* and *C9orf72* expression (Figure 1A & Figure S1A).

**Figure 1.**
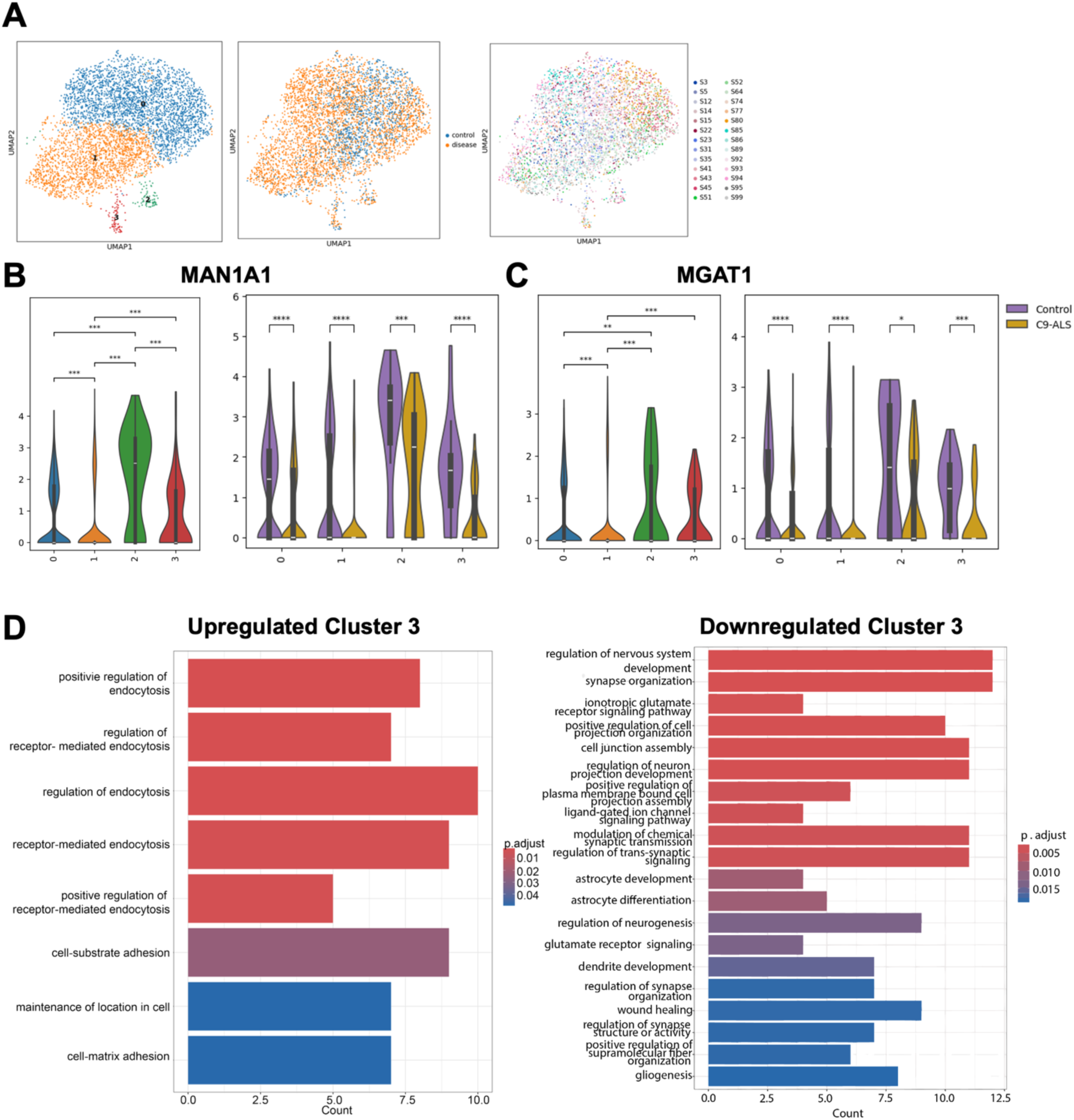
snRNA-Seq from human postmortem frontal cortex indicates altered MAN1A1 expression across microglia clusters and a decrease in C9-nuclei. (A) Uniform manifold approximation and projection (UMAP) visualization of snRNA sequencing-derived microglia sub-cluster (left), representation of disease background (middle) and individual samples (right). n= 13 CTR and n= 10 ALS, and n=6 ALS/FTD tissue. (B) Violin plots of *MAN1A1* expression across microglial clusters and between C9orf72 nuclei and control nuclei. White bar indicates median expression. n= 5 CTR tissue (low PMI), n= 10 ALS tissue and n= 6 ALS/FTD tissue. Statistics were performed with Wilcoxon Rank-Sum with Benjamini-Hochberg correction. *p<0.05, **p<0.01, ***p<0.001, ****p<0.0001 (C) Violin plots of *MGAT1* expression across microglial clusters and between C9orf72 nuclei and control nuclei. White bar indicates mean expression. n= 5 CTR tissue, n= 10 ALS tissue and n= 6 ALS/FTD tissue. Statistics were performed with Wilcoxon Rank-Sum with Benjamini-Hochberg correction. *p<0.05, **p<0.01, ***p<0.001, ****p<0.0001 (D) Pathway enrichment from differentially expressed genes (DEGS) of cluster 3. n= 5 CTR, n= 10 C9-ALS and n= 6 ALS/FTD tissues. Colors indicate the p.adjusted value decreasing in value from blue to red.

Differential expression analysis of combined C9-ALS and C9-ALS/FTD nuclei compared to CTR- nuclei with a low postmortem interval (PMI) revealed one of the most significantly reduced genes across clusters in C9-nuclei was in mannosidase alpha class 1A member 1 (MAN1A1), a Golgi α-mannosidase required for N-glycan maturation, accompanied by increased expression of inflammatory mediators and pathways linked to immune activation (Figure 1B-C & Figure S1B), while cluster 2 showed higher levels of MAN1A1 compared to the other microglia clusters (Figure1B). MAN1A1 is the initiating enzyme in the Golgi to help trim and mature mannose residues^47^. Mannosyl (alpha 1,3-) – glycoprotein beta-1,2-N-acetylglucosaminyltransferase (MGAT1), the sequential enzyme following MAN1A1, is similarly reduced as MAN1A1 in all microglia clusters (Figure 1C). Pathway enrichment demonstrated suppression of neuronal programs, including processes involved in synapse organization, glutamate signaling, and astrocyte differentiation and development, together with activation of processes involved in endocytosis and cell adhesion, suggesting that impaired glycoprotein processing accompanies microglia transition toward a disease-associated state (Figure 1D). To further characterize disruptions in Golgi and glycan metabolism, we performed module scoring using curated gene sets representing relevant biological processes. This analysis revealed a significant increase in nucleotide-sugar biosynthesis and a significant decrease in Golgi calcium homeostasis (q-values; Figure S1C). Several functionally related modules exhibited nominally significant changes but did not meet the false discovery rate (FDR) threshold. Specifically, mannose trimming and nucleotide sugar transport are reduced, while Golgi-to-ER retrograde transport and Golgi ion homeostasis were also indicated with a nominal p-value level (Figure S1C). Although these modules did not remain significant after FDR correction, their close functional relationship to pathways that met the corrected significance threshold suggests that they may reflect broader perturbations in Golgi and glycan-associated processes. Additional studies will be required to determine the extent and biological significance of these alterations. Collectively, these findings identify dysregulated Golgi- associated processes and nucleotide-sugar biosynthesis as previously unrecognized features of microglia in human C9orf72-associated ALS/FTD.

### Secretome profiling identifies defective glycan degradation in C9orf72 microglia

As many glycosylated proteins from the Golgi are directed to the secretory pathway, we wondered whether altered glycosylation signatures extend to microglial secreted proteins, we performed quantitative proteomic analysis of conditioned media (CM) collected from one control and one C9orf72 iPSC-derived microglia line (Figure 2A). Functional enrichment analysis through STRING of microglia conditioned media (MCM) revealed a difference in proteins associated with the lysosome, in addition to alterations in extracellular exosome pathways (Figure 2B-C). Isolating the proteins associated with the *Lysosome* enrichment present in the ACM and performing functional enrichment analysis revealed via KEGG and Molecular Function enrichment terms including *other glycan degradation* (Figure 2D) and *hydrolase activity* (Figure 2E) as alternately enriched in C9-ACM. The proteins identified as part of these terms from the *Lysosome* related protein population are highlighted in a protein-protein interaction network (Figure 2F), and from this another mannosidase enzyme, MAN2B1, was identified in both; which is reduced 2-fold in ALS/FTD MCM (Figure 2D-G). Further investigation with more power is needed to determine a statistically significant difference of MAN2B1 expression between control and disease. This finding is consistent with impaired glycan turnover and hydrolysis of mannose residues in the lysosome (Figure 2D-G)^48^. These observations support the findings from the snRNA seq analysis and suggest that C9orf72 microglia exhibit widespread defects in glycoprotein processing and secretion. Together, these data indicate that abnormalities in glycan metabolism are reflected not only within microglia but also throughout the extracellular signaling environment they create through their secretome.

**Figure 2.**
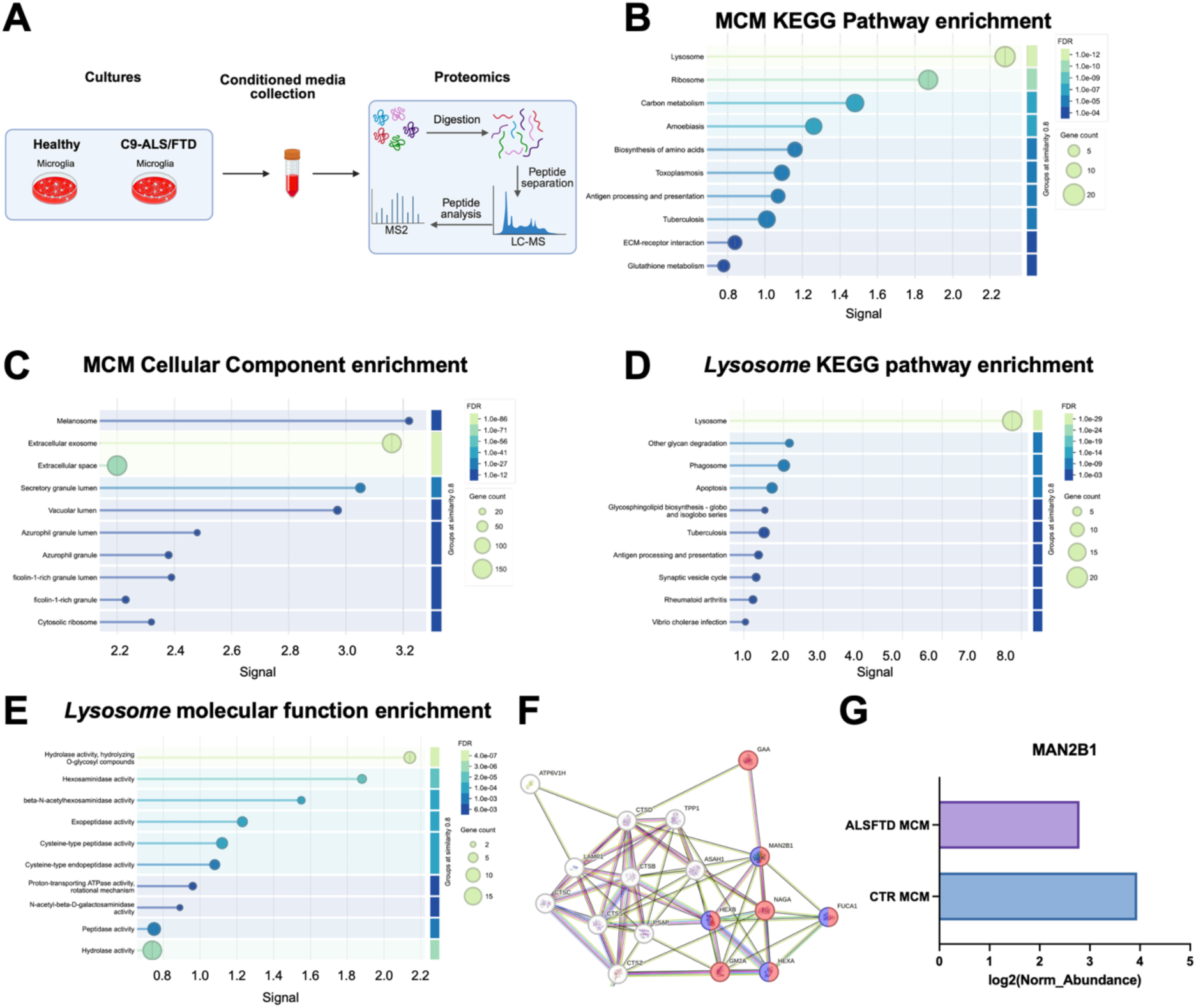
STRING enrichment analysis of MCM proteome identifies lysosomal disruption to glycan degradation in C9-MCM. (A) Schematic overview of experimental workflow regarding microglia conditioned media (MCM) collection. (B-C) Pathway enrichment analyses of differentially abundant proteins identified in MCM (KEGG and GO cellular component). (D) Pathway enrichment analysis (KEGG) of the *Lysosome* term proteins identified in (B). (E) Gene Ontology enrichment (molecular function) of the *Lysosome* term proteins identified in (B). (F) Protein-protein interaction network from *Lysosome* highlighting interactions among proteins involved in *other glycan* highlighted in red, and *hydrolase activity, hydrolyzing O-glycosyl compounds* highlighted in purple. Nodes represent individual proteins, and connecting edges indicate known or predicted functional associations. (G) Bar graph indicating log2 of normalized abundance of MAN2B1. Proteins with |log2| >0.5 assessed from n= 1 CTR and 1 ALS/FTD iPSC MG line. Lollipop diagrams display enriched biological pathways, with enrichment signal plotted on the x-axis, bubble size representing the number of proteins associated with each pathway, and color indicating the false discovery rate (FDR)-adjusted significance.

### C9orf72 iPSC astrocytes secretome reveals impaired ER-Golgi protein processing and reduced MAN1A1 levels

Knowing that microglia are surveying cells taking cues from the environment to inform their function and role^49^ and because astrocytes strongly influence microglial states, we next examined the astrocyte secretome using quantitative proteomics, with the goal of later using this media to treat iPSC microglia. Astrocyte conditioned media from diseased C9orf72 iPSC cortical astrocyte conditioned media (dACM) were enriched for extracellular matrix (ECM) organization, complement terms, while localization of enrichment was identified to be associated with the endoplasmic reticulum lumen and largely differences in extracellular components, strongly suggesting altered extracellular signaling and vesicle composition in C9-ACM when compared to control cell line ACM (cACM) (Figure 3A- B). Assessing only proteins decreased in dACM compared to cACM indicated selective depletion of proteins involved in neurodegeneration, including *ALS*-*proteasome*, and *protein processing in the endoplasmic reticulum (ER)* (Figure 3C- D). Notably, *protein processing in the ER* included asparagine N-linked glycosylation- the type of glycosylation affected by mannosidases like MAN1A1- while MAN1A1 itself appeared within this network alongside valosin-containing protein (VCP), a known ALS associated risk factor (Figure 3F-H)^50^. These observations suggest that astrocytes are an important source of altered glycoprotein processing signals that may influence neighboring microglia. To further understand how this might impact the molecular profile and functional consequences resulting from a microglia-astrocyte crosstalk, the ACM from both healthy and C9orf72 cortical astrocytes was used to acutely treat iPSC MG.

**Figure 3.**
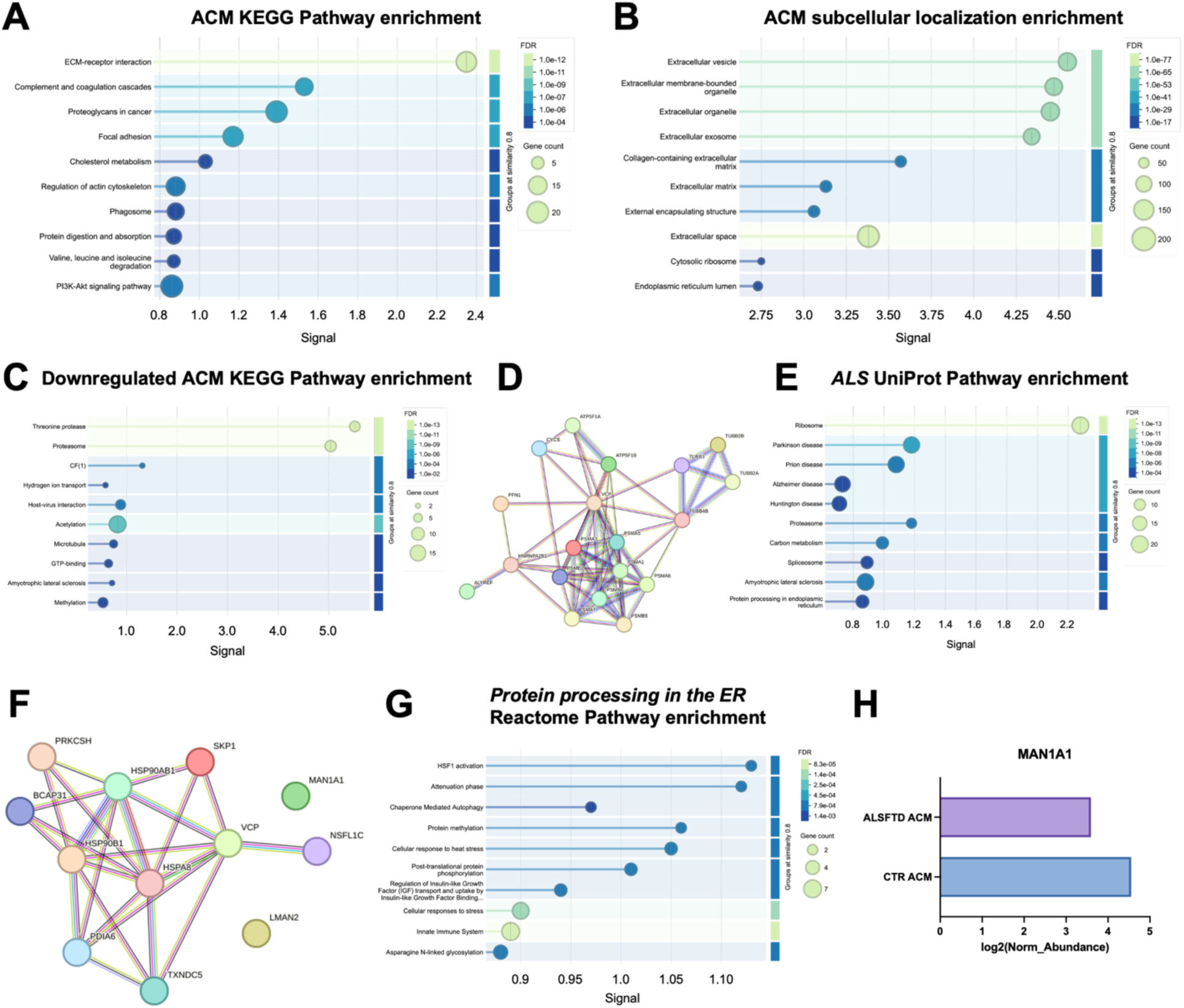
Proteomic pathway enrichment and protein network analyses of ACM reveal disrupted proteostasis of the proteasome and ER protein processing. (A-B) Functional enrichment analysis (KEGG) of differentially abundant proteins identified in diseased astrocyte conditioned media (dACM). Lollipop diagrams display enriched biological pathways, with enrichment signal plotted on the x-axis, bubble size representing the number of proteins associated with each pathway, and color indicating the false discovery rate (FDR)- adjusted significance. Proteins with |log2| >0.5 used for analysis of n= 1CTR cell line and n= 1 ALS/FTD cell line. (C) Pathway enrichment analysis (KEGG) from proteins decreased in dACM compared to cACM, highlighting pathways associated with protein turnover and cellular homeostasis. Bubble size, color, and enrichment signal are represented as described in (A). Proteins with log2< −0.5 used for analysis of n=1 CTR and n=1 ALS/FTD cell line. (D) Protein-protein interaction network of the *ALS* pathway proteins identified in (C). (E) Pathway enrichment analysis (UniProt keywords) of *ALS* pathway proteins shown in (D). (F) Protein-protein interaction network of the *Protein processing in the ER* pathway identified in (C). (G) Pathway enrichment analysis (reactome) of the *Protein processing in the ER* pathway proteins shown in (F). Nodes represent individual proteins, and connecting edges indicate known or predicted functional associations. H) Bar graph indicating log2 of normalized abundance of MAN1A1 abundance.

### Healthy astrocyte secretome promotes Golgi remodeling and MAN1A2 expression in microglia

To determine how astrocytes regulate microglial proteostasis, healthy iPSC-derived microglia were conditioned with ACM, followed by transcriptomic and proteomic profiling of microglia cell pellets. iPSC cortical astrocytes from non-neurological controls (c) were cultured and matured for 95 days, and astrocyte conditioned media (ACM) was collected over two days (Figure S2A). Meanwhile, iPSC microglia were cultured to maturity and primed with LPS for 2 hours before being exposed to healthy control ACM (LPScACM) for 6 hours (Figure 4A-B)^51^. Bulk transcriptomic and proteomic analyses were performed on microglial cell pellets to determine molecular shifts on both the RNA and protein level when compared to standard microglia maturation media fed microglia cultures (MGMAT). Transcriptomic analysis revealed a statistically significant separation between MGMAT and ACM-treated samples, while this separation was less distinct at the protein level (Figure S2B-C). Performing STRING analysis on proteins with abundance (|logFC| > 0.5) in CTR-MG treated with cACM revealed that exposure to healthy ACM induced coordinated activation of RNA and protein binding, altered exosomes and mitochondrial metabolism, suggesting increased metabolic processing in microglia exposed to ACM (Figure 4C). Markov cluster algorithm (MCL) analysis of these same proteins revealed 48 clusters with three or more proteins. Increased spliceosome engagement, and alternative RNA splicing, in addition to increased translation were revealed as the top clusters (Figure 4D-E). Notably, reactome pathway enrichment revealed increases in proteins related to transport to the Golgi, Intra-Golgi and Golgi-to-ER trafficking in a subsequent cluster within the top 15 identified clusters (Figure 4F). ANOVA analysis showed significance of MAN1A2 across groups (p= 0.025) but was unable to identify which groups contribute to this variance, suggesting more power is needed to isolate specific group contributions. (Figure 4F-G). These data suggest that astrocytes under physiological conditions promote a biosynthetically active microglial state characterized by enhanced Golgi processing and protein homeostasis.

**Figure 4.**
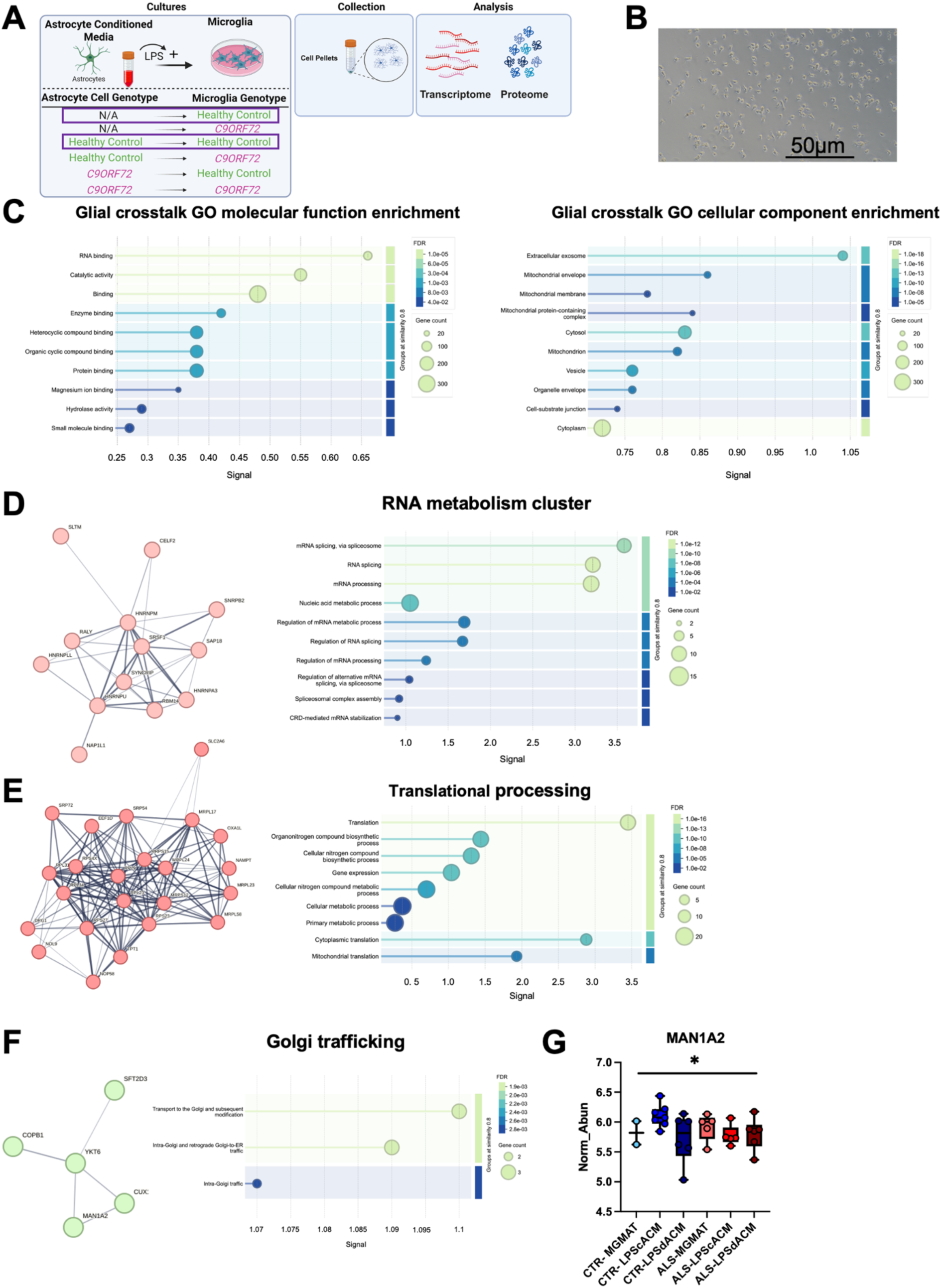
Alterations in RNA processing and protein synthesis revealed in microglia exposed to healthy ACM. (A) Schematic overview of the experimental design. Human healthy control iPSC-MG were primed with LPS before being exposed to healthy control astrocyte conditioned media (ACM). Following treatment, MG cell pellets were collected for transcriptomic and proteomic analyses. Purple boxes indicate treatments utilized in enrichment analysis. (B) Representative phase-contrast image of microglia following exposure to ACM. Scale bar, 50mm. (C) Functional enrichment analyses of molecular changes identified through proteomic profiling highlighting molecular function and cellular component gene ontology enrichment. Lollipop diagrams display enriched biological pathways, with enrichment signal plotted on the x-axis, bubble size representing the number of associated proteins, and color indicating false discovery rate (FDR)-adjusted significance. Proteins with |log2FC| > 0.5 were used for analysis. n= 7 CTR and n =7 ALS/FTD cell lines. (D) Protein-protein interaction network from top cluster from STRING Markov clustering (MCL) derived from differentially abundant proteins (DAPs) in CTR-MG treated with cACM. Highlighting RNA processing and metabolism, and *spliceosome* function, consistent with alterations in RNA regulatory processes. An inflation parameter of 3 was used. E) Protein-protein interaction network from second STRING Markov clustering (MCL) derived from differentially abundant proteins in CTR-MG treated with cACM identifies biological process enrichment of proteins involved in *translation*. An inflation parameter of 3 was used. F) STRING MCL clustering of DAPs from CTR-MG treated with cACM indicates *transport to the Golgi* within top 15 clusters identified. An inflation parameter of 3 was used. Nodes represent individual proteins, and connecting edges indicate known or predicted functional associations. G) Bar graph showing normalized MAN1A2 protein abundance in MG pellets. n= 7 CTR and 7 ALS/FTD cell lines. ANOVA with Tukey’s post-hoc analysis * p< 0.05, **p< 0.01, ***p<0.001, ****p<0.0001. Error bars indicate standard error of the mean.

### Diseased astrocyte-microglia crosstalk fails to support microglial Golgi homeostasis and instead induce inflammatory remodeling

We wondered whether C9orf72 astrocytes’ secretome (diseased ACM; dACM) differentially affects the function of healthy microglia and C9orf72 microglia (Figure 5A). Phase contrast imaging did not reveal noticeable morphological changes in C9orf72 iPSC-microglia cultures (Figure 5B). Control and C9orf72 microglia were primed with LPS for 2 hours, followed by a 6 hour treatment with either cACM or dACM (LPScACM; LPSdACM). STRING network analysis of protein abundance |logFC| > 0.05 of cell lysates from control microglia treated with LPScACM and C9orf72 microglia treated with LPSdACM revealed alterations in terms associated with exosomes and other vesicular compartments, in addition to terms relating to the extracellular matrix (ECM) (Figure S3A). UniProt keyword associations and Molecular Function enrichment further indicate altered *proteoglycan binding* and a link to *glycogen storage disease* components (Figure 5C &D). Within these two enrichment assessments, several disease-associated signaling molecules were significantly (ANOVA with Tukey’s post-hoc analysis) altered when C9orf72 microglia were treated with the C9orf72 astrocyte secretome, highlighted below. Consistent with acquisition of a reactive microglial phenotype, APOE – with multiple allelic forms which lead to altered protein function^52,53^ and a known genetic risk factor for developing Alzheimer’s Disease (AD) - was identified in the *glycogen storage disease* network together with THBS1; *glycogen storage disease* proteins highlighted in red (Figure 5C)^54^. APOE is downregulated while THBS1 is elevated when C9orf72 microglia were exposed to LPSdACM compared to control microglia exposed to LPScACM (Figure 5E). Of note, THBS1 is also upregulated in control microglia exposed to LPSdACM (Figure 5E). In the *proteoglycan binding* term (associated proteins highlighted in red) both acid alpha- glucosidase (GAA) and phosphoglucomutase 1 (PGM1) protein abundances were significantly altered in response to C9orf72 ACM treatment (Figure 5D & 5F). GAA is a critical lysosomal protein responsible for the breakdown of glycogen into glucose and is shown to be downregulated in the disease context, while PGM1 is upregulated (Figure 5F). Overactivation of this enzyme leads to an imbalance of cytokine production^55^. Importantly, performing a t-test analysis was able to reveal C9orf72 ACM reduced MAN1A2 levels (p= 0.047) in C9orf72 microglia compared to CTR-MG treated with cACM (Figure 5G), suggesting that disease astrocytes fail to maintain Golgi glycoprotein processing in microglia while simultaneously promoting extracellular matrix remodeling and inflammatory activation.

**Figure 5.**
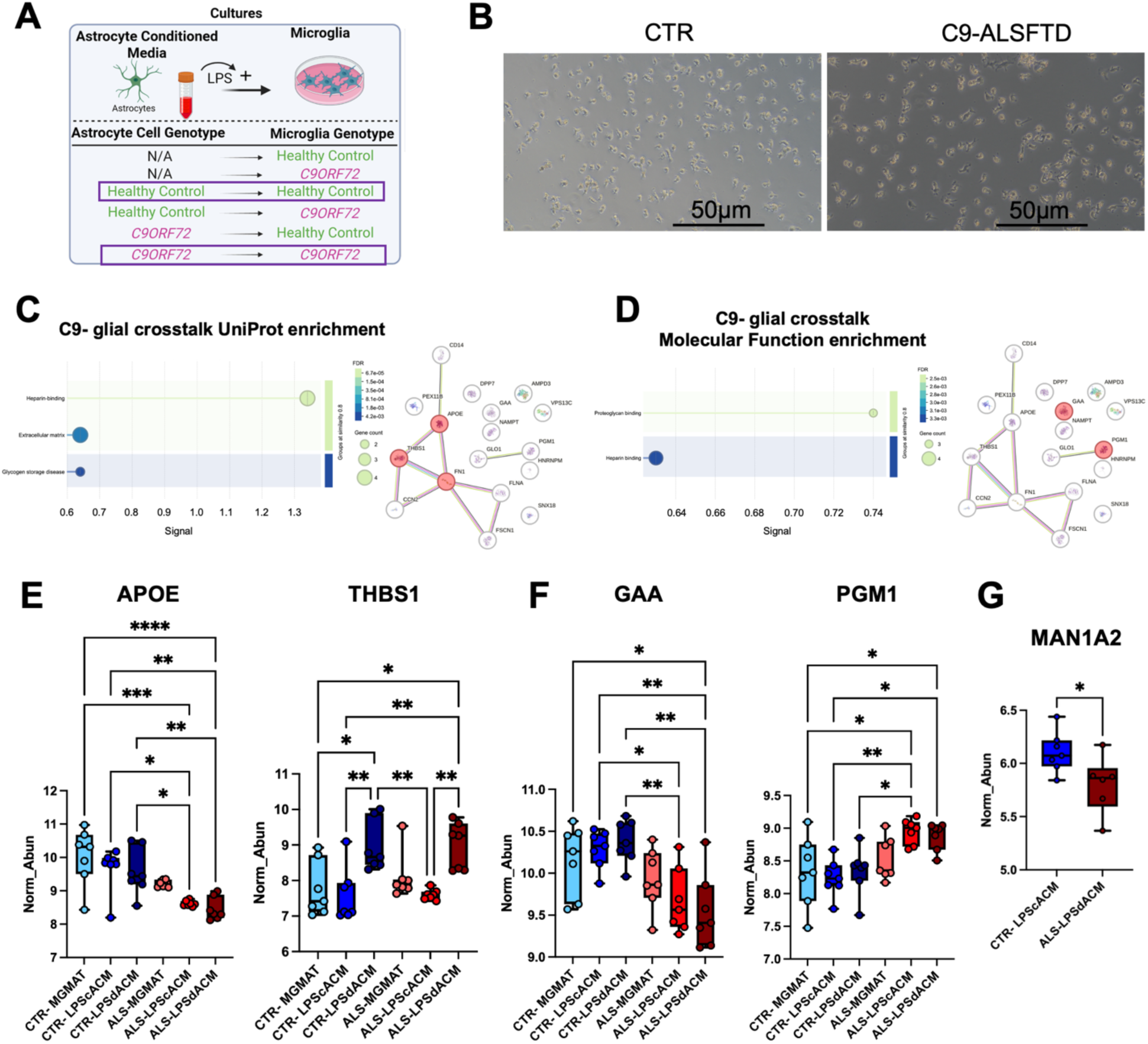
C9-glial-crosstalk indicates disruption to energy maintenance and neuroinflammatory pathways. (A) Schematic overview of the experimental design. Human healthy control and C9orf72 (C9) iPSC-MG were primed with LPS followed by exposure to healthy control or C9 astrocyte conditioned media (LPScACM or LPSdACM, respectively). Groups assessed in the following enrichment analysis are indicated in the purple boxes. (B) Representative phase-contrast image of control (CTR) and C9-ALS/FTD microglia at D57. Scale bar, 50μm. (C) Left: Gene ontology molecular function enrichment analysis (STRING) highlighting *glycogen-storage disease*-related annotations among DAPs identified in C9-MG treated with LPSdACM. Enrichment signal is plotted on the x-axis, bubble size represents the number of associated proteins, and color indicates false discovery rate (FDR)-adjusted significance. Right: Protein-protein interaction network highlighting proteins associated with *glycogen storage disease*, highlighted in red, including fibronectin (FN1), thrombospondin-1 (THBS1) and apolipoprotein E (APOE). n= 7 CTR and n= 7 ALS/FTD cell lines. (D) Left: Functional enrichment analysis identifying *proteoglycan-binding* and *heparin-binding* activities among proteins altered in C9-MG treated with LPSdACM. Enrichment signal is plotted on the x-axis, bubble size represents the number of associated proteins, and color indicates false discovery rate (FDR)-adjusted significance. Right: Protein-protein interaction network highlighting proteins associated with *proteoglycan binding* ontology, including acid alpha-glucosidase (GAA) and phosphoglucomutase 1 (PGM1) –highlighted in red. n= 7 CTR and 7ALS/FTD cell lines. Nodes represent individual proteins, and connecting edges indicate known or predicted functional associations. (E-F) Quantitative comparisons of the indicated protein abundances in MG pellets across experimental groups. Individual points represent biological replicates, with bars showing group means. One-way ANOVA with Tukey’s post-hoc test. * p< 0.05, **p< 0.01, ***p<0.001, ****p<0.0001. Error bars indicate standard error of the mean. G) Boxplot of MAN1A2 protein abundance in MG pellets comparing healthy MG exposed to LPScACM to C9-MG exposed to LPSdACM. Welch’s t-test * p< 0.05 n= 7 CTR and n=7 ALS/FTD cell lines. error bars indicate standard error of the mean.

### Integrated multi-omics identifies inflammatory and redox dysregulation during C9orf72 glial-crosstalk

To determine whether altered astrocyte–microglia crosstalk promotes inflammatory activation in C9orf72 ALS/FTD, we quantified cytokine release from microglia following ACM exposure and integrated these findings with transcriptomic and proteomic analyses. Multiplex cytokine profiling (Meso Scale Discovery, MSD) revealed altered secretion of several microglial inflammatory mediators including IL-12p40, IL-2, IL-6, TNFα, and IL-1α (Figure 6A; Figure S3B-F). Notably, interferon gamma (IFNγ), and VEGFA were significantly increased in C9orf72 microglia, while VEGFA was also elevated in control microglia exposed to C9orf72 ACM (Figure 6B). Together with the increased THBS1 abundance in C9orf72 microglia following disease ACM exposure (Figures 5E,6B), these findings identify altered VEGFA/THBS1-associated signaling as a potential component of disease-related glial-crosstalk.

**Figure 6.**
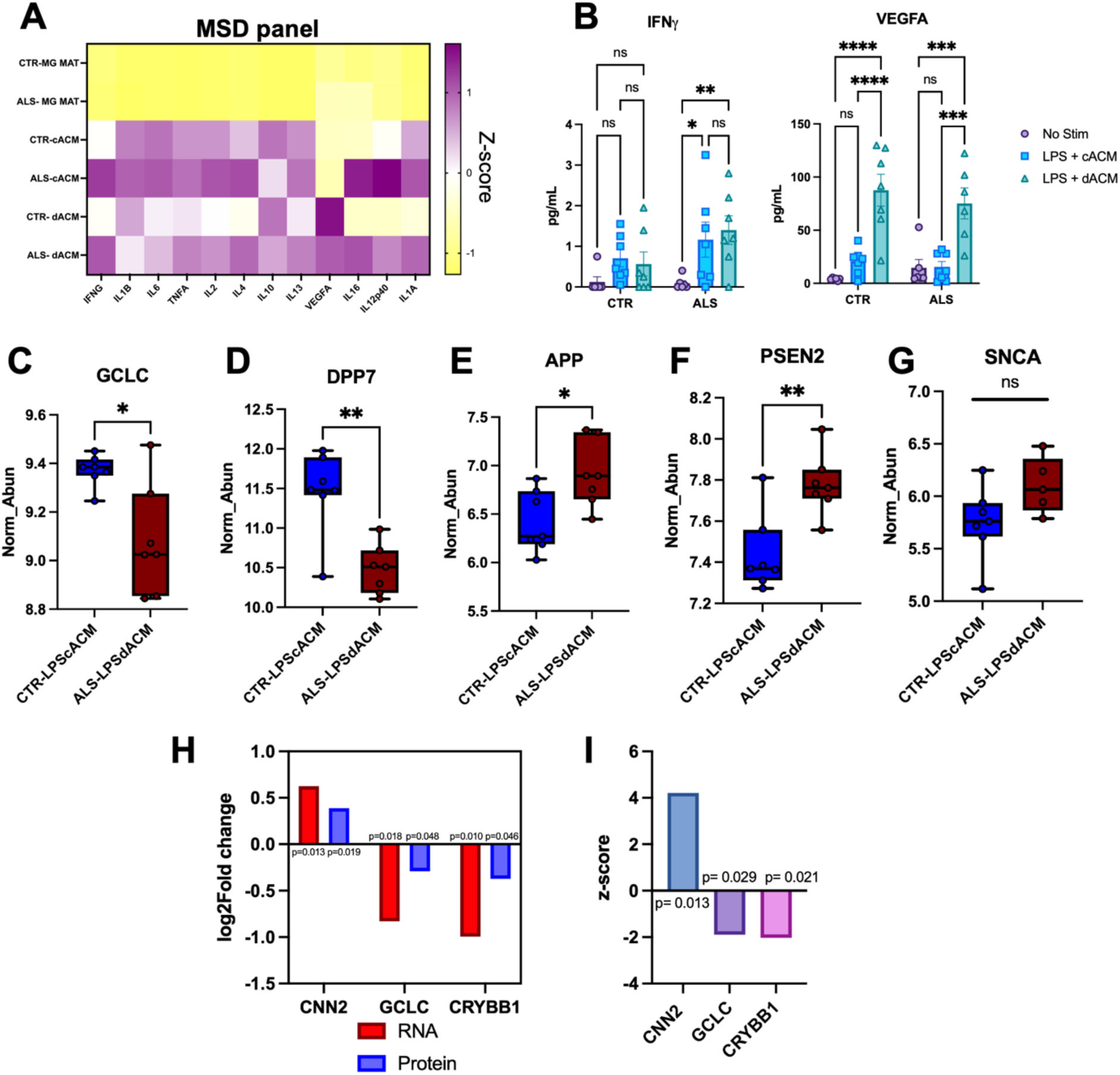
Astrocyte-conditioned media alters Inflammatory responses and reveals changes in disease-associated molecular pathways in microglia. (A) Heatmap showing relative levels of inflammatory mediators measured using a multiplex Meso Scale Discovery (MSD) assay in conditioned media collected from healthy control (CTR) and C9orf72 ALS/FTD (ALS) microglia following exposure to varying ACM conditions. Color intensity represents standardized abundance (Z-score), with yellow indicating lower and purple indicating higher levels. (B) Quantification of IFN**γ** and VEGFA in conditioned media from CTR and ALS microglia under unstimulated conditions or following ACM treatment. n= 6 CTR and 6 C9 samples. Two-way ANOVA. (C-G) Quantitative proteomic analysis comparing normalized protein abundance in CTR-MG exposed to cACM (CTR-LPScACM) and C9 MG exposed to C9 ACM (ALS-LPSdACM). Box plots show normalized abundance of glutamate-cysteine ligase catalytic subunit (GCLC; C), dipeptidyl peptidase 7 (DPP7; D), amyloid precursor protein (APP; E), presenilin 2 (PSEN2; F), and α-synuclein (SNCA; G). n=7 CTR and n=7 C9 samples. Error bars indicate standard error of the mean. Welch’s t test. H) Comparison of transcriptomic (red) and proteomic (blue) Log2 fold changes in calponin 2 (CNN2), GCLC, and crystallin beta B1 (CRYBB1) in MG cell pellets. T-test with Bonferroni method p-value indicating C9-MG treated with LPSdACM compared to CTR-MG treated with LPScACM. I) Standardized expression scores (Z-score) for CNN2, GCLC, and CRYBB1, highlighting differences in the relative magnitude and direction of molecular changes across these targets. Adjusted p-value with Benjamini-Hochberg method indicating changes in C9-MG treated with dACM compared to CTR-MG treated with cACM. * p< 0.05, **p< 0.01, ***p<0.001, ****p<0.0001.

This inflammatory response was accompanied by changes in genes and proteins associated with cellular stress and neurodegeneration. Comparison of LPS-primed control microglia treated with cACM and LPS-primed C9orf72 microglia treated with dACM identified glutamate-cysteine ligase catalytic subunit (GCLC), a rate-limiting enzyme in glutathione synthesis, was reduced at both the transcript and protein levels in C9orf72 microglia exposed to dACM, consistent with altered redox (Figure S3G, Figure 6C).^56,57^. Proteomic analysis further identified reduced dipeptidyl peptidase 7 (DPP7), and altered abundance of the neurodegeneration-associated proteins amyloid precursor protein (APP), and presenilin 2 (PSEN2), whereas *α* -synuclein (SNCA) was unchanged. (Figure 6D–6G).

Integrated analysis generating z-scores from combined protein abundance and RNA transcript quantification and performing statistics on these scored identified three significantly dysregulated at both levels: GCLC, CNN2, and CRYBB1 (Figure 6H). GCLC and CRYBB1 were significantly decreased (p= 0.029 and p=0.021, respectively) from z-score analysis, whereas CNN2 (p= 0.013) was significantly increased in combined analysis (Figure 6I). These molecules link several of the processes emerging from our earlier analyses (Figure S3B, Figure 6C). These molecules link several of the processes emerging from our earlier analyses: GCLC is linked with cellular redox homeostasis and inflammation, CRYBB1 has been associated with microglia homeostasis and lysosomal/phagocytic function, and CNN2 helps modulate macrophage motility, and has been identified as a target of VCP^58–64^. STRING network analysis further linked CNN2 to IL-12/JAK- STAT signaling (Figure S3H) and superoxide dismutase 1 (SOD1), a gene whose variants are associated with familial ALS (Figure S3H)^65,66^.

Collectively, these data demonstrate that C9orf72 astrocyte–microglia crosstalk induces a coordinated microglial response characterized by inflammatory activation, impaired redox homeostasis, and altered neurodegeneration-associated proteins. Together with the impaired MAN1A1/MAN1A2 abnormalities identified above, these findings further connect altered glycan and secretory pathway biology with inflammatory reprogramming of C9orf72 microglia.

### Interactor analysis identifies APOE-, THBS1-, and VCP- associated signaling networks

To further define signaling between astrocytes and microglia, we performed Ingenuity pathway- interactor analysis used to predict upstream regulators by integrating ACM and microglia cell pellet proteomic and transcriptomic datasets, restricting the analysis to nervous system-, microglia-, and astrocyte- relevant molecules. This identified interactions between 18 proteins/genes detected in microglia and multiple molecules from ACM-derived proteins/genes (Figure 7A). Of these interactions networks involving APOE and transforming growth factor beta 1 (TGF-β1) were identified (Figure 7A). APOE is the only significantly differentially abundant microglial derived molecule within these networks, as noted previously (Figure 5E, Figure 7B). The identified interactors for APOE include SNCA, APP and ciliary neurotrophic factor (CNTF) (Figure 7A). Two SNCA isoforms were identified in ACM, isoform 2-4 was detected only in disease ACM, whereas isoform 2-5 was present in both control and disease ACM (Figure 7C-D). APP abundance was comparable between conditions (Figure 7E), while CNTF was higher in dACM. CNTF is a component of the cortical astrocyte culture media therefore this abundance cannot be unequivocally attributed to astrocyte secretion (Figure 7F).

**Figure 7.**
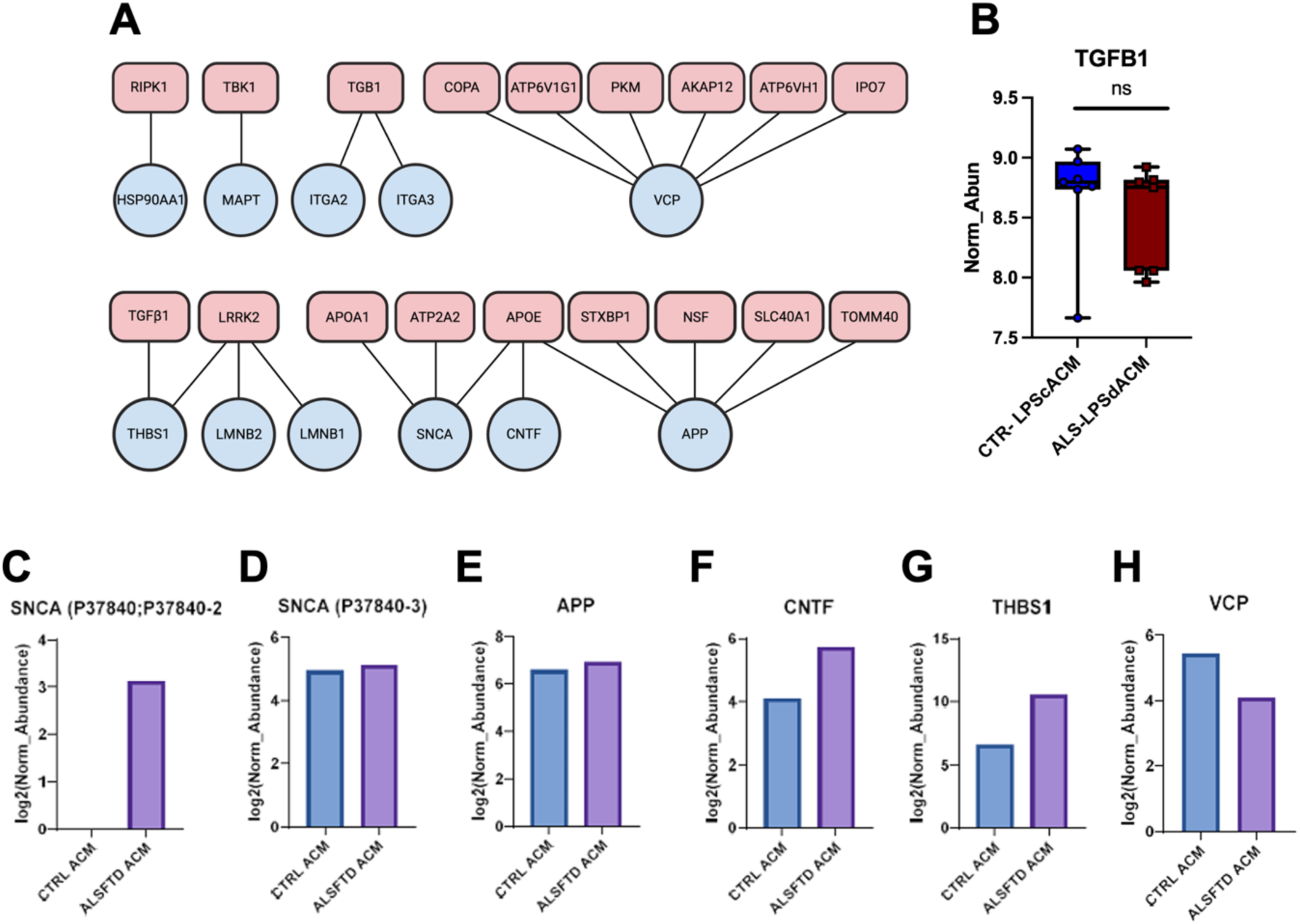
Protein interaction analysis reveals disease-associated proteins and altered abundance of APOE, THBS1, and VCP in C9 astrocyte conditioned media. (A) Ingenuity pathway analysis (IPA, Qiagen) identified interactions of proteins present in microglia cell pellets (pink squoval) with proteins present in astrocyte conditioned media (blue circles). (B) Quantification of transforming growth factor beta 1 (TGFB1) protein abundance in control microglia exposed to control ACM (CTR-LPScACM) and C9orf72 ALS/FTD microglia exposed to C9orf72 ALS/FTD ACM (ALS-LPSdACM). Box plot show normalized protein abundance, with individual points representing biological replicates. No statistically significant difference was detected between these two conditions (ns = not significant). n= 7 CTR and 7 C9 samples. Significance was calculated by Welch’s t-test. (C-H) Relative abundance of two SNCA protein isoforms (C+D), APP (E), CNTF (F), THBS1 (G) and VCP (H) in control (CTR) versus C9orf72 ALS/FTD (ALS/FTD) astrocyte conditioned media. Values are expressed as log2-transformed normalized protein abundance. n= 1 CTR and 1 ALS/FTD cell line.

TGF-β1 abundance in microglia was unchanged, while its interacting ligand THBS1 is increased in MG and showed higher levels in the C9orf72 ACM (Figure 7G), further implicating altered THBS1-associated signaling in C9orf72 glial-crosstalk.

Interactor analysis additionally identified reduced VCP abundance in disease ACM, and 6 interacting proteins identified in MG including alpha- COP (COPA); pyruvate kinase muscle (PKM), ATP6V1G1, and A-kinase anchoring protein 12 (AKAP12) (Figure 7A, 7H). Thus, independent network analysis converged on proteins associated with ALS/FTD and vesicular, Golgi, and lysosomal biology, reinforcing the pathway-level alterations identified by the preceding multi-omic analysis (Figure 6 and Figure 7).

To assess the disease relevance of these findings, we examined 15 identified targets from iPSC disease microglia exposed to dACM when compared to control microglia exposed to cACM, discussed throughout this work, in the postmortem snRNA-seq dataset. The only significantly differentially expressed genes meeting FDR thresholding from the list of 15 targets were PKM (increase) and MAN1A1 (decrease) in C9-ALS and ALS/FTD nuclei. Both of which were consistent with iPSC glial-crosstalk modeling findings. Other genes were assessed on trends relative to controls, *AKAP12*, *VCP*, *ATP6V1G1*, *CNN2*, *APP*, and *THBS1* were increased in both C9-ALS and C9-ALS/FTD microglia, whereas *APOE*, *CRYBB1*, and *GAA* were decreased in both disease groups (Figure S4A). Directional changes between iPSC model and postmortem microglia were concordant for 10 out of 15 targets in C9-ALS and 8 of 15 targets in C9-ALS/FTD (Figure S4B), providing independent support for disease-relevant pathways identified in the human iPSC glial-crosstalk model.

## Discussion

Largely acknowledged as a source of pathology in ALS, neuroinflammation‘s key mechanisms from glial-crosstalk have been unexplored until recently^3^. While cytokine, chemokine, and trophic factor signaling are key consequences of glial-crosstalk, studies on the intracellular consequences of these mechanisms tend to stop at individual molecule activation and inhibition^20,67–69^. This work highlights glycosyl metabolism as a novel influential process in the neuroinflammatory network of microglia and astrocytes in C9orf72 ALS/FTD by identifying dysregulation of glycosyl metabolism of the ER, Golgi, and lysosome. Glycosylation of proteins is the major role of the Golgi apparatus; where O-linked glycosylation occurs and N-linked proteins from the ER are matured^39^. Glycoproteins are typically directed from the Golgi apparatus to the lysosome or the plasma membrane, and are commonly secreted proteins^39,70^. They are major components of cell-to-cell signaling, extracellular matrix, and lysosomal membranes^39^. Additionally, glycolipids are critical for nerve insulation, immune defense, and cell signaling^71–73^. Given these vital roles, alterations in glycosylation and glycosyl metabolism can have broad impacts on cellular function.

We identified a reduction in the initiating enzyme of N-glycosylation in the Golgi, MAN1A1 in addition to the sequential enzyme MGAT1, in microglia from human postmortem tissue of ALS patients (Figure 1B & 1C). Notably, enrichment analysis revealed functional disruption to neuronal homeostasis via synaptic organization, glutamate signaling, and astrocyte differentiation and development (Figure 1D). This mannosidase enzymatic process was reduced in an iPSC model of microglial, where levels of the developmentally enriched form of MAN1A1, MAN1A2, were increased in control microglia treated with LPScACM compared to C9orf72 microglia treated with LPSdACM. Loss of function of this protein has been linked to lysosomal storage disorders in addition to an association with autism spectrum disorder, intellectual disability, and dementia^48,74^.

Secretome profiling from both microglia and astrocytes further highlighted glycosyl metabolism, identifying altered enrichment in C9orf72 for lysosomal degradation and hydrolyzation specifically of glycans, in addition to extracellular aberrations (Figure 2B-E). The reduced levels of MAN2B1 secreted by diseased microglia suggest probable disruption to the degradation of complex sugars (Figure 2G)^48^. Accumulation of complex sugars can overwhelm the endolysosomal pathway, resulting in the secretion of improperly processed products^75^. The astrocytic secretome implicates similar disruptions to the extracellular matrix and vesicle signaling. Protein processing in the ER revealed specific N-glycan metabolism. Specifically, MAN1A1 was reduced in this network, also linking MAN1A1 to ALS associated protein, VCP (Figure 3F). VCP is implicated in Golgi maintenance^76^. Lysosomal disruption has been identified as a hallmark of ALS pathology and this work identifies a potential source of this pathology^77,78^. Degradation of glycoproteins and glycolipids within the lysosome is critical for recycling the building blocks of cellular signaling and is a key source of fuel when energy needs arise^79–81^. Disruption to these turnover processes can lead to storage diseases^79–82^. This supports a potential convergence of ALS with lysosomal storage disorders. Secretome findings need to be explored further due to limited sample size.

Given the additional association of failed turnover with cellular energy^82^, it is unsurprising that processes linked to oxidative stress pathways also emerged. THBS1 is a glycoprotein, known to regulate cell adhesion, migration and angiogenesis in addition to playing a role in innate immune activation^83^. ER stress has been shown to lead to THBS1 presentation on the surface of cells, but can also activate protective stress responses including nuclear factor erythroid 2-related factor (NRF2) signaling^84^. NRF2 signaling has been proposed as a therapeutic target for ALS^84,85^. THBS1 and VEGF are part of a signaling cascade that influence angiogenesis, immune regulation, and cell proliferation and migration^86^. These two signals were both increased in microglia in the presence of dACM, suggesting astrocytic dependent activation of this signaling cascade. In addition, interactor analysis identified TGFβ1-THBS1 signaling as a potential stimulant for this increase (Figure 7A). It has been noted in others’ work that silencing THBS1 leads to decreased VEGFA^83^. VEGFA release is also influenced by the ECM and can be released with altered metalloproteinase-9^87^. Interestingly, VEGF mRNA has been shown to be regulated by another ALS/FTD associated protein ELAVL3^88–90^. Additionally, VEGFA is a high-mannose N- linked glycoprotein meaning it requires glycosyl metabolism^91^. If glycosyl metabolism is dysregulated in C9-ALS/FTD, as indicated by our results, the glycosylation of both THBS1 and VEGFA may be impacted which would potentially affect the function and turnover of these proteins. Additional oxidative stress cues were identified including GCLC. GCLC is critical for glutathione production, a master antioxidant, and also linked to NRF2 signaling^57,92–96^. Conditional knockout of GCLC in mouse neurons induced mitochondrial damage and neurodegeneration^56^. Moreover, increases in cytokine and chemokine levels, complement proteins, enhanced phagocytosis, and the activation of pyroptosis via gasdermin is observed with GCLC deficiency^56,57^. This data couples identified disruptions from glial-crosstalk in ALS/FTD with the oxidative stress observed in disease.

Proteoglycan binding was identified in network analysis indicating a reduction in GAA from C9orf72 glial-crosstalk. Deficiencies in GAA lead to the development of Pompe’s disease, which leads to damage of the heart and skeletal muscles in addition to affecting spinal cord motoneurons and cortical tissues^97^, likely due to its role in processing glycogen. This enzyme is created as a precursor glycoprotein and requires processing into its mature active form^98^. PGM1 was also identified from the proteoglycan binding network. It provides substrates required for glycoprotein and carbohydrate synthesis, and is associated with Congenital Disorder of Glycosylation, Type 1t^99^. Patients display a LOF in this gene leading to developmental deformities, growth delays, and myopathy^100^. In the context of myeloid cells, PGM1 activation increases metabolic processes to meet needs for early immune activation and stress response^55,101^. This further coalesces glycosyl metabolism, metabolic demand, and oxidative stress, while also indicating links to neuromuscular pathways involved in ALS.

While C9orf72 pathology, including TDP-43 dysfunction, primarily results in ALS/FTD progression, TDP-43 pathology is also recognized as a mechanism of AD pathology^102,103^. This work further highlights this association, identifying APOE, APP and PSEN2 disruptions, while interactor analysis defined signaling cascades present amongst these factors. Additionally, DPP7 was identified via proteomics to be downregulated (Figure 6C). It has been shown that reduced Dpp expression can accelerate neurodegeneration via innate immune response in drosophila flies^18^. DPP7 is known for being a quiescent lymphocyte marker, and is normally located in vesicles and at the Golgi^62^, however, knowledge of its function is limited. Other DPP proteins, such as DPP4 or DPP9, are known to regulate synaptic plasticity and alter LTP and memory function^104,105^. Notably, consistent with our findings of altered Golgi processing and glycoproteins, these processes have been recently implicated in proteomic analysis from CSF and plasma of AD and FTD patients^106^. Neuroinflammation and microglia function have been identified as key mechanisms in AD, with APOE being an essential modifier of Aβ plaques derived from alternative cleavage of APP^54,107^. While APOE is most notably described by its allelic associated risk factor, the consequence of the APOE ɛ4 allele is critically altered protein function promoting a pro- inflammatory state, impairing phagocytic function, and disrupting lipid metabolism^52^. Together, this suggests that C9orf72 leads to an altered microglia state that is shared between ALS, FTD and AD. A potential cause of this molecular shift is glycosylation state and the handling of glycosylated products.

This work also highlighted VCP in both network analysis (Figure 3E) and the interactor analysis (Figure 7A). Interactor analysis revealed a reduction in secretion of VCP from C9orf72 astrocytes, and identified multiple proteins associated with its signaling found in MG including COPA, ATP6V1G1, PKM, and AKAP12. COPA is associated with a syndrome, COPA syndrome, a rare immune disease^108^. ATP6V1G1 is V-type proton ATPase subunit which has functions in lysosome, exosome, and synaptic vesicle acidification as well as iron homeostasis^109^. It has previously been linked to ALS via UBQLN2 studies in which UBQLN2 fails to bind ATP6V1G1 leading to reduced levels resulting in the inability to break down products via autophagic pathways^110^. Interestingly, pyruvate kinase muscle (PKM), a rate-limiting enzyme in glycolysis, was identified as a receptor for VCP (Figure 7A). PKM is noted to be upregulated in spinal neurons from ALS models, and pharmacological inhibition of this enzyme aggravated pathology in ALS mouse models^111^. It is suggested that PKM plays an early protective role, alleviating mitochondrial stress^111^. Another notable receptor of VCP is A-kinase anchoring protein 12 (AKAP12) (Figure 7A). AKAP12 is a scaffolding protein that binds proteins like PKC near the cell membrane, of which PRKCB was found upregulated in the snRNA-Seq data (Figure 1D)^112^. Others have noted changes in PKC of ALS patients, with various clinical trials aimed at targeting protein kinases (Figure 1D, Figure 7A)^9,113,114^. Together this data shows how dysregulation of VCP signaling, along with its interactors, can have wide reaching effects including on immune reaction, oxidative stress, ECM and extracellular signaling in addition to the major roles VCP plays in the cytoplasm including autophagy, Golgi structure, mitochondrial performance, and ER morphogenesis^115^. These signaling molecules were validated to have differential expression in disease from human postmortem tissue, and also highlighted differences in expression between C9-ALS and C9- ALS/FTD indicating potential use in differentiating C9orf72 disease progression into a more cognitive or motor disease (Figure S4A).

Finally, this work highlights overlap between proteins associated with subtypes of ALS including C9orf72, VCP, SOD1, and UBQLN2 forms. VCP was identified in the network analysis from ACM, linked with MAN1A1 (Figure 3F). Additionally, the multi-omic analysis identified CNN2, which has also been identified as a target of VCP (Figure 6H). Furthermore, IL-12 STRING network projections link IL-12, CNN2, and SOD1 (Figure S3H). Together this implicates shared mechanisms across disease subtypes, and highlights how the findings from this work can impact the disease spectrum. The results from this work provide a framework to investigate potential therapeutic targets and biomarkers of disease. Further detailed studies aimed at identifying specific altered glycosylation and glycosyl metabolism will be needed to explore potential biomarkers and targets further.

## Limitations

Several limitations of the experimental design should be considered when interpreting these findings. First, although C9orf72 HRE is the most common genetic cause of ALS and FTD and is strongly associated TDP-43 pathology in human disease, overt TDP-43 pathology is not recapitulated in the iPSC models of MG or CA here. Thus, these models are best suited to interrogate C9-dependent glial alterations and mechanisms that may occur upstream of, or independently from, TDP-43 pathology. Second, our crosstalk studies were designed to capture relatively early glial responses and were limited to single time points and acute conditioned media exposures. The duration and concentration of conditioned media treatments were also limited by abundance of iPSC microglia resulting from differentiation, which were within anticipated numbers. While this design enabled the identification of early cellular and molecular responses to reciprocal glial signaling, exploring additional timepoints, longer exposures, or alternative exposure strategies such as concentrated media may provide further insight into disease mechanisms. Our integration of these findings with previously published snRNA-seq data introduces additional limitations inherent to that dataset, including a relatively small number of control samples due to the need to match metrics of postmortem intervals (PMI) between control and disease cases. Down sampling was used to address this limitation, nevertheless, these factors may limit the ability to resolve less abundant cell states or more subtle disease-associated transcriptional changes. Finally, while the convergence of our findings on Golgi and lysosomal pathways and proteins involved in glycosyl metabolism supports altered protein processing as a potential feature of C9orf72 glial dysfunction, the current studies do not directly define changes in specific glycosylation. Targeted glycomics analyses will therefore be necessary to determine the specific protein and lipid glycosylation changes associated with C9orf72 microglia-astrocyte communication.

## Resource availability

### Lead contact

Request for further information and resources should be directed to and will be fulfilled by the lead contact, Rita Sattler.

### Materials availability

This study did not generate new unique reagents/materials. The iPSC lines used in this study have been generated previously and are available for distribution from the Cedars-Sinai Biomanufacturing Center (https://biomanufacturing.cedars-sinai.org/).

### Data and code availability

- Single-nuclei RNA-seq data from Gittings et al 2023^36^ had been deposited at https://www.synapse.org/Synapse:syn45351388/wiki/622682
- RNA sequencing transcriptomics data has been deposited in the GEO Submission (GSE348042) and Mass spectrometry proteomics data has been deposited to the ProteomeXchange Consortium via the PRIDE partner repository with the dataset identifier (PXD085088).
- Any additional information required to reanalyze the data reported in this paper is available from the lead contact upon request.

## Acknowledgements

The authors would like to thank the Integrated Mass Spectrometry Shared Resource at TGEN for the processing of protein samples and the Collaborative Sequencing Center at TGEN for processing of RNA samples. This work was supported by the NIH-NINDS (grants 1F31NS139617-01A1 (B.N.O) and R01NS120331 (R.G.S and K.VK.J); Muscular Dystrophy

Association (MDA 1063968; R.G.S); Robert Packard Center for ALS Research; Barrow Neurological Foundation (B.N.O).

## Author contributions

B.N.O and R.G.S designed the experiments, wrote, and edited the manuscript. Oversight of experimental execution was given by R.G.S, K.VK.J and P.P. B.N.O performed *in vitro* experiments with assistance from L.M.G, K.R.P, and J.L,. E.A, B.M, K.VK.J performed analysis for snRNA-seq data. K.G.M, N.H, B.L, R.S, M.M, and P.P performed mass spectrometry and proteomics analysis. A.B, S.S, I.S.P and M.H performed RNA sequencing and analysis for iPSC bulk-transcriptomics and integrated multi-omics. Input on manuscript was provided by all co- authors. B.N.O: Briana N Ondatje; L.M.G: Lauren M Gittings; K.R.P: Kim R Preller; EA: Eric Alsop; K.G.M: Krystine Garcia-Mansfield, N.H: Nate Hansen; B.L: Brooke Lovell; R.S: Ritin Sharma; I.S.P: Ignazio S Piras; M.M: Melissa Martinez; B.M: Bessie Meechoovet; A.B: Anna Bonfitto; S.S: Serena Song; J.L: Jennifer Levy; M.H: Matthew Huentelman; P.P: Patrick Pirrotte; K.VK.J: Kendall van Keuren- Jensen; R.G.S: Rita Sattler

## Declaration of interests

The authors do not have any disclosures.

## Declaration of generative AI and AI-assisted technologies in the manuscript preparation process

Generative AI was utilized to help organize and curate Golgi and glycosylation module gene lists.

## Supplemental information titles and legends

Document S1. Supplemental Figures 1-4

## STAR Methods

### Experimental model and study participants

#### Cell lines

iPSC lines used in our studies were purchased from Cedars Sinai-induced pluripotent stem cell core https://www.cedars-sinai.edu/research/areas/biomanufacturing/ipsc.html and are part of the Answer ALS cohort ^116^. A Certificate of Analysis (COA) for these lines is available upon request.

#### Cluster cell type annotation and subclustering

Leiden clusters were annotated using established marker genes from cortex samples, as previously described^36^ by analyzing expression levels of marker genes found in each cluster. The following marker gene list was used: AQP4 and GFAP (astrocytes), EPSA1 and CLDN5 (endothelial cells), RBFOX3 (neurons), SATB2, SLC17A7 and NRGN (excitatory neurons), GAD1 and GAD2 (inhibitory neurons), ADAM28 and APBB1IP (microglia), RNF220 and ST18 (oligogendrocytes), PDGFRA and SMOC1 (oligodendrocyte progenitor cells; OPCs). Microglia and astrocyte subclusters were determined further subclustering just these cluster using the Leiden algorithm at a cluster resolution of 0.2, resulting in four microglia and four astrocyte subclusters.

#### SnRNA-seq differential expression analysis

Astrocyte and microglia snRNA-seq data was pseudobulked using the sc.get.aggregate() function in scanpy on patientID, diseaseGroup, subcluster and sex. Pseudobulk counts were loaded in R and analyzed using DESeq2 with diseaseGroup and sex as covariates in the model. CSV files of the DESeq2 results were retained for pathway analysis.

#### Pathway analysis of snRNA-seq

Pathway analysis was performed using clusterProfiler (enrichGO function v4.18.4) against Gene Ontology in R Studio. Genes which were upregulated were identified with logFC > 0.3 and p_adj.loc > 0.05. Genes with a logFC < −0.3 and p_adj_loc > 0.05 were considered downregulated.

#### Gene module analysis

Manually curated gene sets representing twenty-five modules were scored per cell in Scanpy using scanpy.tl.score_genes with default parameters. Per-cell scores were aggregated to donor- level values by taking the median across all cells of a given cell type within each donor and donor– cell type combinations with fewer than 10 cells were excluded. Astrocytes and microglia were analyzed independently. Within each cell type, module scores were compared between control samples (n = 5) and disease samples (n = 16; C9-ALS and C9-ALS/FTD combined) using two- sided Welch’s t-tests, with Benjamini–Hochberg correction applied across all twenty-five modules within each cell type. Heatmaps display the median module score per group, with per-module p- values and FDR-corrected p-values annotated alongside each row.

#### Microglia cell differentiation

iPSCs were differentiated into microglia following the protocol by Moser et al 2021^117^ with slight modifications consistent with Lorenzini et al 2023^27,118,119^, described here. iPSCs were grown in 6 well plates coated in 167μg/mL Matrigel in mTeSR plus at 37°C and with 5% O_2_ until they reached 80-90% confluency. Cells were passaged using a StemPro EZPassage Disposable Stem Cell Passaging Tool (#23181010, Life Technologies) and plated on matrigel in mTeSR plus. Cells were maintained over the next 7 days, removing differentiating cells and colonies growing within touching distance of another colony. Once wells had roughly 10–12 colonies measuring 1.0 mm in diameter on average, differentiation was started using stage 1 media, consisting of 80 ng/ml bone morphogenetic protein 4 (BMP4) in mTeSR plus. Media was changed daily for 4 days, cells were then switched to stage 2 media, consisting of 25 ng/ml basal fibroblast growth factor (bFGF), 100 ng/ml stem cell factor (SCF) and 80 ng/ml vascular endothelial growth factor (VEGF) in StemPro-34 serum free medium. Two days later, cells were switched to stage 3 media, consisting of StemPro-34 supplemented with 50 ng/ml SCF, 50 ng/ml interleukin-3, 5 ng/ml thrombopoietin (TPO), 50 ng/mL macrophage colony stimulating factor (M-CSF) and 50 ng/mL FMS-related tyrosine kinase 3 ligand (Flt3). Media was changed 4 days later and replaced with stage 3 media until day 12, when cells were switched to stage 4 media, consisting of 50 ng/ml M-CSF, 50 ng/ml Flt3 ligand, and 25 ng/ml granulocyte-macrophage colony stimulating factor (GM-CSF) in StemPro-34 medium. Three days later, 1mL of stage 4 media was added to the wells. Four days later, the floating cells in the supernatant were collected, pelleted, resuspended in fresh stage 5 media consisting of 1x MEM NEAA, 1x pen/strep, 1x Glutamax, 1x

B27, 1x N2, 1x insulin transferrin, and 5μg/mL insulin, 400uM 1-thioglycerol, 25ng/mL M-CSF, 100ng/mL IL-34, and 50ng/mL TGF-beta in DMEM/F12, and placed in two new Matrigel coated plates, with even distribution among the wells. Four days later cells were fed with an additional 1mL of stage 5 media. Original plates were fed with 1mL of stage 4 media to continue maintaining the hematopoietic stem cell population. Original plates were fed with an additional 1mL of stage 4 media four days later. These collection feeds were performed once per week to begin microglial fating, and original plates were fed with stage 4 media for up to 6 weeks. On D52, cells were collected and plated into final plate formats in RPMI-1640 medium supplemented with 1x GlutaMAX, 10 ng/ml M-CSF, and 100 ng/ml interleukin −34 and 50ng/mL TGF-beta, 100ng/mL CD200, and 100ng/mL CX3CL1. Stimulation with ACM was performed five days later. 50k cells for 24 wells and 150k for 6wells.

For stimulation, cells treated with LPS had media replaced with 100ng/mL LPS in MGMAT without small molecules for 2 hours, cells without LPS had MGMAT without small molecules. After 2 hours, cells were fed with their respective astrocyte conditioned media for 6 hours. Cells without ACM were fed MGMAT without small molecules. Cells were enzymatically harvested by accutase treatment for 10 min at 37°C, collected and pelleted, and stored at −80°C until RNA extraction or protein preparation.

#### Astrocyte cell differentiation

80-90% confluent iPSCs were differentiated into an initial neural progenitor cell (NPC) state following Shi et al 2019^120^. Briefly, cells were dissociated with accutase and seeded in a monolayer into a Matrigel coated well with 2mL of mTeSR plus. The following day media was replaced with 3mL of neuronal induction media (NIM) consisting of 1x B27, 1x N2, 1x MEM NEAA, 1x Glutamax, 1x Pen/Dtrep, combined with 2.5μg/mL insulin, beta-mercaptoethanol, 10μM SB431542, LDN193189 in a 1:1 DMEM/F12 to Neurobasal media. NIM was replaced every day for 12 days. Following the 12 days, cells were split using StemPro EZPassage with each well of a 6 well placed into two new 6 well wells. 3mL of neuronal maturation media was used with Rock inhibitor (1:2000). Neuronal maturation media consists of 1:1 DMEM/F12 and neurobasal with 1x B27, 1x N2, 1x MEM NEAA, 1x Glutamax, 1x Pen/Strep, and 2.5μg/mL insulin, 100μM 2-mercaptoethanol plus the addition of added growth factor to the NMM for four days 20ng/mL bFGF. After four days of NMM + FGF, media was replaced with NMM without growth factors every day until D22 of differentiation. Cells were then banked using a control rate freezer and placed in liquid nitrogen. Frozen NPCs were thawed into Matrigel coated plates into NPC media containing DMEM/F12 with 1x B27, 1x Pen/Strep, 10ng/mL EGF, 10ng/mL bFGF, and 2ng/mL heparin. Cells were fed with NPC for 7 days until they were split into Matrigel coated T75 flasks using accutase dissociation. Dissociated cells were resuspended into stage 1 astrocyte (S1A) media containing DMEM/F12, 1x B27, 1x Pen/Strep, and 20ng/mL LIF. Cells were fed every three days with S1A media for fourteen days, with passaging occurring once per week into new Matrigel coated flasks, size dependent on cell line growth. After this time, cells are passaged into stage 2 astrocyte media (S2A) containing DMEM/F12, 1x B27, 1x Pen/Strep, 20ng/mL LIF, 20ng/mL CNTF, and 20ng/mL BMP4. Cells were fed S2A every three days until D60 of differentiation when they were banked using a control rate freezer and placed into liquid nitrogen. Banked astrocytes were thawed into Matrigel coated flasks for further maturation in S2A + growth factors 20ng/mL EGF, 20ng/mL FGF, and 2ng/mL heparin. Cells were fed every other day with S2A + growth factors until D93 of differentiation, after cells were fed with S2A every other day. D95 astrocyte conditioned media was collected for use as treatment.

#### Astrocyte conditioned media collection

Media was collected after 2 days from the D95 astrocyte flask, spun at 300rcf for 10min to remove cellular debris, supernatant was collected and placed in a new conical, and frozen in −80 until further use.

#### Microglia conditioned media collection

Microglia cultures were collected, spun down at 300rcf for 10min, placed in fresh media and conditioned for 8 hours total. Media was collected from microglia plates, spun at 300rcf for 10min to remove cellular debris, supernatant was collected and placed in a new conical, and frozen in −80 until further use.

#### Bulk-transcriptomic sample prep

Qiagen RNeasy Micro RNA kits were used for RNA isolation of microglia cell pellets. RNA was assayed for quantity and quality respectively by Qubit (cat# Q32855/Q10211, Thermo Fisher Scientific) and Tapestation (cat# 5067-5579/5067-5576, Agilent Technologies). Libraries were created according to manufacturer protocol using KAPA RNA HyperPrep Kit with RiboErase (HMR) (cat# KK8561, Roche). After pooling, libraries were sequenced on a NovaSeqX instrument (Illumina).

#### Bulk-proteomics sample prep cell pellets

Samples were dried by speed-vac and lysed in 25uL of 8M urea lysis buffer. Samples were digested using an in-solution trypsin digestion. Protein concentration was quantified using the Pierce BCA Protein Assay Kit (ThermoScientific, Cat# 23225).

Mass spectrometry data were acquired on a NanoElute2 liquid chromatography system (Bruker Daltonic) coupled to timsTOF Ultra mass spectrometer (Bruker Daltonic). 50ng of peptides per sample were directly loaded on a 25cm C18 column (PepSep ULTRA, 75uM ID, 1.5 μM particle size) and separated by a 30 min Reverse Phase chromatography gradient at a flow-rate of 250nL/min. Data acquisition was performed in diaPASEF mode over a mass tnge of 300-1000m/z and a mobility range (1/K0) of 0.64-1.20 with a cycle time of 0.96s. Capillary voltage was at 1600V and dry gas temperature was at 200 degrees Celsius. Accumulation and ramp time was 100ms with 100% duty cycle.

#### Bulk-proteomics sample prep conditioned media

125μg of protein was digested using an in-solution trypsin digestion. Peptides were quantified using the Pierce BCA assay (ThermoScientific, Cat# 23225).

Mass spectrometry data was acquired on a NanoElute2 liquid chromatography system (Bruker) coupled to timsTOF Ultra mas spectrometer (Bruker). 50ng of peptides per sample were directly loaded on a 25cm C18 column (PepSep ULTRA, 75μM ID, 1.7 uM particle size) and separated by a 30 min Reverse Phase chromatography gradient at a flow-rate of 250nL/min. Data acquisition was performed in diaPASEF mode over a mass range of 300-1000m/z and a mobility range (1/K0) of 0.64-1.20 with a cycle time of 0.96s. Capillary voltage was at 1600V and dry gas temperature was at 200 degrees Celsius. Accumulation and ramp time was 100ms with 100% duty cycle.

#### Protein Identification

Mass spectrometry data from cell pellets were searched in Spectronaut (v19.2) in directDIA mode against predicted Human Swissprot/UniprotKB protein sequencing database (downloaded May 2020). Trypsin theoretical digestion was used allowing for up to 2 missed cleavages. Cysteine carbamidomethylation was set as a fixed modification. Variable modifications were protein N-term acetylation and methionine oxidation. Precursors, peptides, and protein groups were filtered for FDR <0.01. Normalization and imputation were not performed within Spectronaut. All other default settings were retained. Variance stabilization normalization (VSN) was used to normalize abundances. Data filtering and PCA plots were performed in R v4.4.0.

#### Data analysis proteomics

Initial significance was assessed using a one-way ANOVA followed by Tukey post-hoc to correct for multiple hypothesis testing for pairwise comparisons. Significance threshold set at p-value < 0.05 (Tukey adjusted). Two valid data points were required for Tukey post-hoc calculation. Proteins with logFC > 0.5 or < −0.5 were processed in STRING Network analysis.

#### STRING enrichment analysis

Enrichment analysis was performed using the STRING v12 online interface using default settings and filters. Proteins from proteomic analysis showing a log fold change >0.5 or < −0.5 were used for the analysis. Markov cluster Algorithm analysis, finding clusters based on stochastic flow, was used with an inflation parameter of 3. Pathways and GO enrichment also performed within STRING. Terms were grouped by similarity ≥ 0.8 and sorted by signal. Proteins identified in enrichment analyses were analyzed in R studio v4.5.3, where boxplots were generated.

#### Ingenuity Pathway Analysis

To analyze microglia receptor interactions, the microglia proteomics dataset was imported into Ingenuity Pathway Analysis (IPA; QIAGEN) and filtered for human nervous system receptor proteins^121^. Filtering parameters were restricted to human species (species = Homo sapiens), relevant tissue and cell types (tissues & cell lines = nervous system, astrocyte, microglia, neurons), and specified receptor node types (node types = G-protein coupled receptor, growth factor, ion channel, kinase, ligand dependent nuclear receptor, transmembrane receptor, transporter) yielding a dataset of 916 proteins. The IPA build tool was used to add direct downstream interactors to the receptor network using the same tissue and species parameters. Interactors were restricted to those directly downstream of the receptors. The ACM proteomics data was overlayed onto the built network to identify direct interactors of microglia that were identified in the ACM dataset. Subsequently, the ACM RNA-seq data was overlayed onto the network to confirm transcript level identification for the microglia interactors.

#### Integrated multi-omic analysis

Microglial bulk RNA-seq reads were processed with nf-core/rnaseq v3.14.0 (PMID: 40731283) using STAR–Salmon, the GRCh38 human primary assembly, and Ensembl annotation release 113. Samples with fewer than 10 million total reads or less than 80% uniquely mapped reads were excluded. Low expressed genes with 72 or fewer counts summed across samples were removed. Differential expression was analyzed with DESeq2 (PMID: 25516281) using a design that included sex and the combined microglial-background/treatment group. An omnibus likelihood-ratio test compared this design with a sex-only model. We integrated differential RNA-sequencing and proteomic results by mapping gene symbols to human Ensembl gene IDs using biomaRt and matching the two datasets by Ensembl ID. Protein entries containing multiple gene symbols were split before mapping. Within each comparison, genes were prioritized when both the RNA and protein results had Bonferroni adjusted P < 0.05 across the predefined comparisons, irrespective of the direction of change.

#### Meso Scale Discovery analysis

We performed a V-PLEX Neuroinflammation Panel 1 Human Kit assay (Meso Scale Discovery, Cat #: K15210D) on cell supernatants to assess the state of inflammation following the provided protocol. We looked for a large panel of both pro- and anti-inflammatory cytokines and chemokines, which included GM-CSF, IL-1alpha, IL-5, IL-7, IL-12/IL-23p40, IL-15, IL-16, IL-17A, THF-beta, VEGF-A (MSD, Cytokine Panel 1 human kits Cat#: 15050D), and Eotaxin, MIP-1beta, Eotaxin-3, TARC, IP-10, MIP-1alpha, MCP-1, MDC, and MCP-4 (MSD, Chemokine Panel 1 Gen. B Kits, Cat# K15705). Briefly, plates were washed with wash buffer, followed by the addition of undiluted samples, or prepared calibrators. Samples were left shaking for 2 hours. Plates were then washed with wash buffer, and detection antibody was added to each well as directed shaking for 2 hours. Plates were washed again and read buffer T was added to cytokine panel plates, while Gold read buffer was added to chemokine plates, as directed. Plates were read on a MSD Quickplex SQ 120. Analysis was initially performed within MSD Discovery Workbench, extracted, and analyzed and plotted into bar plots and heatmap in GraphPad Prism v11. Z-scores were generated by subtracting the mean of the molecule expression divided by the standard deviation. Bar plots were generated using a full-fit model Two-Way ANOVA analysis with multiple comparisons comparing the means of each group, error bars display standard error of the mean.

#### Figure design and visualization

Illustrations depicting timelines, experimental paradigms, and visual summaries were created using Biorender.com. Graphs, heatmaps, and other plots were generated with R studio ggplot2 v4.0.3 or enhanced Volcano v1.28.2 and GraphPad Prism v11 and assembled in Adobe Illustrator.

## Notes

https://www.synapse.org/Synapse:syn45351388/wiki/622682

### Competing Interest Statement

The authors have declared no competing interest.

