## Supplemental Figures 1-4 for "Disrupted mannosidase enzymes in C9orf72 ALS/FTD"

A

B

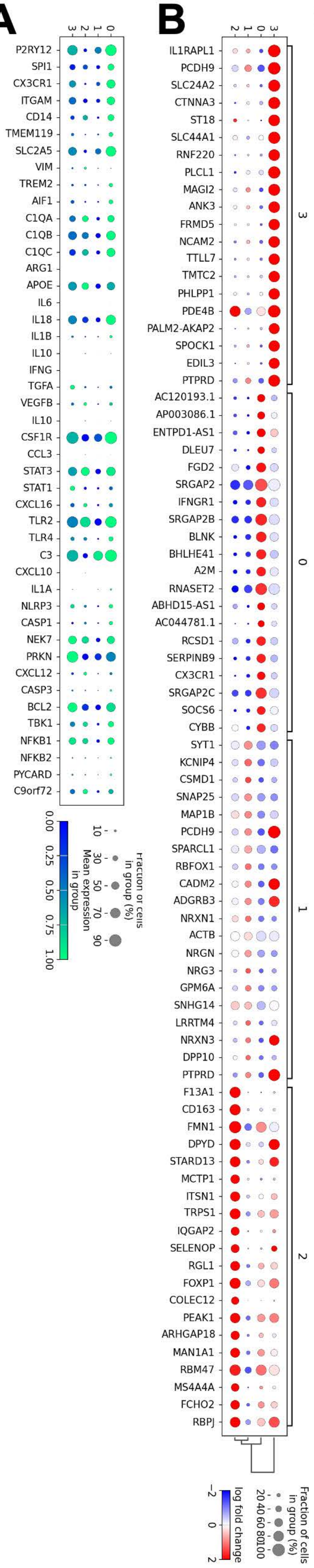


C

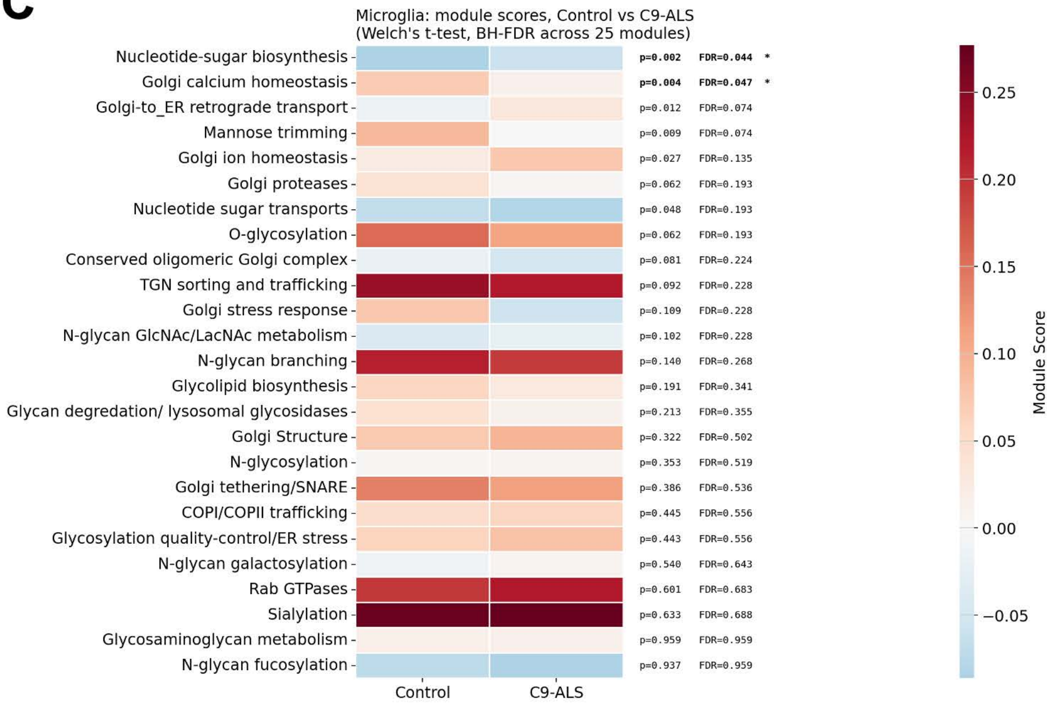


**Figure S1. snRNA-Seq cluster profiles of microglia from human postmortem tissue**

A) Dot plot of selected homeostatic and inflammatory markers in microglia. Colors indicate mean expression while the size of dot represents fraction of cells in group.

B) Dot plot of cluster defining genes in microglia. Colors indicate log2 fold change while the size of dot represents fraction of cells in group.

C) Module scoring of Golgi and glycosylation related processes in microglia nuclei. Bold statistics indicate significant modules padj < 0.5. n= 6 CTR, and 10 C9-ALS and ALS/FTD tissues.

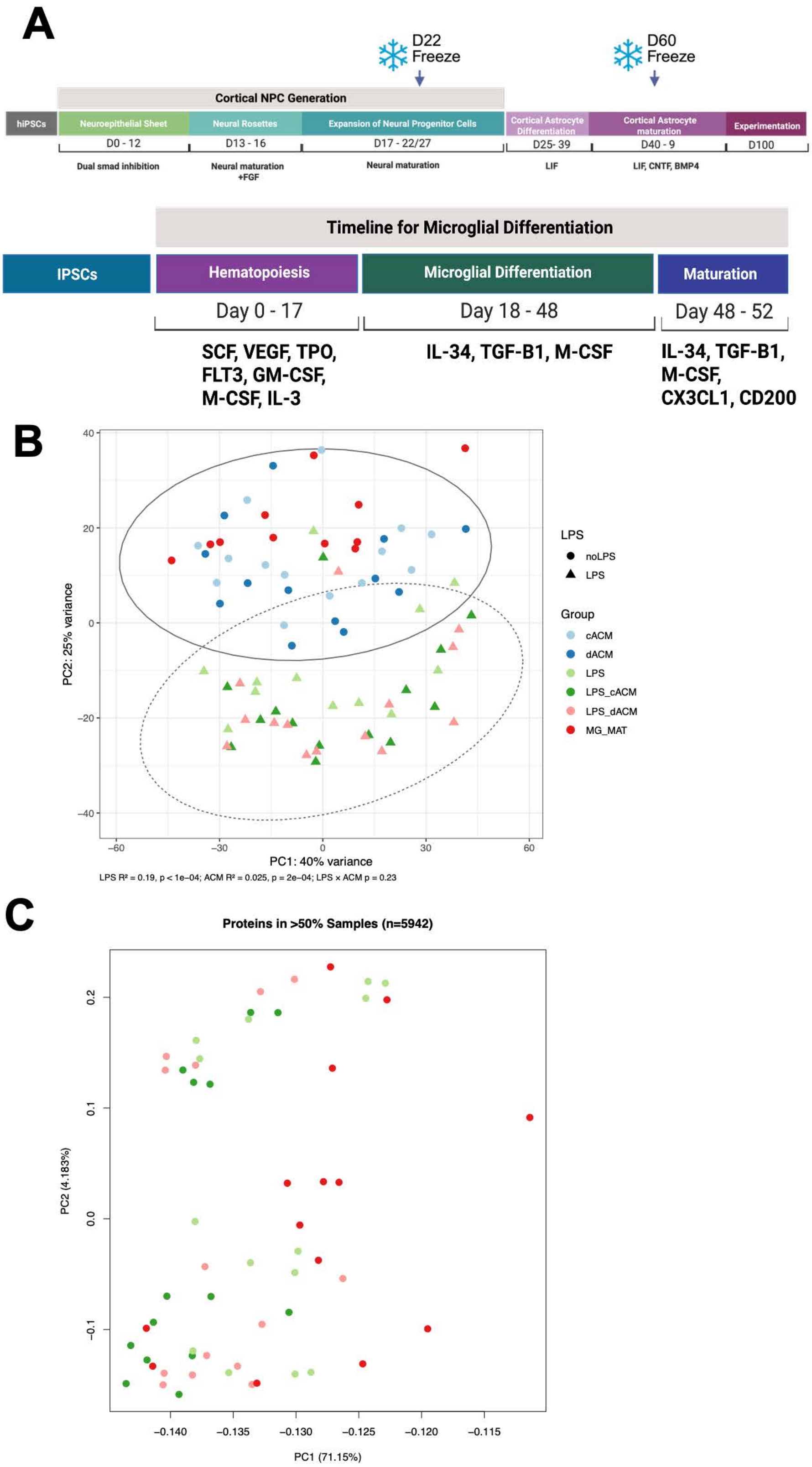


**Figure S2. Transcriptomic and proteomic analysis of ACM treated MG**
A) Diagram of CA (top) and MG (bottom) differentiations.
B) PCA plot of transcriptomics from MG cell pellets. PERMNOVA analysis. LPS $R^2$= 0.19, p< 1e-04; ACM $R^2$= 0.025, p= 3e-04; LPS x ACM p=0.21.
C) PCA plot of proteomics from MG cell pellets.

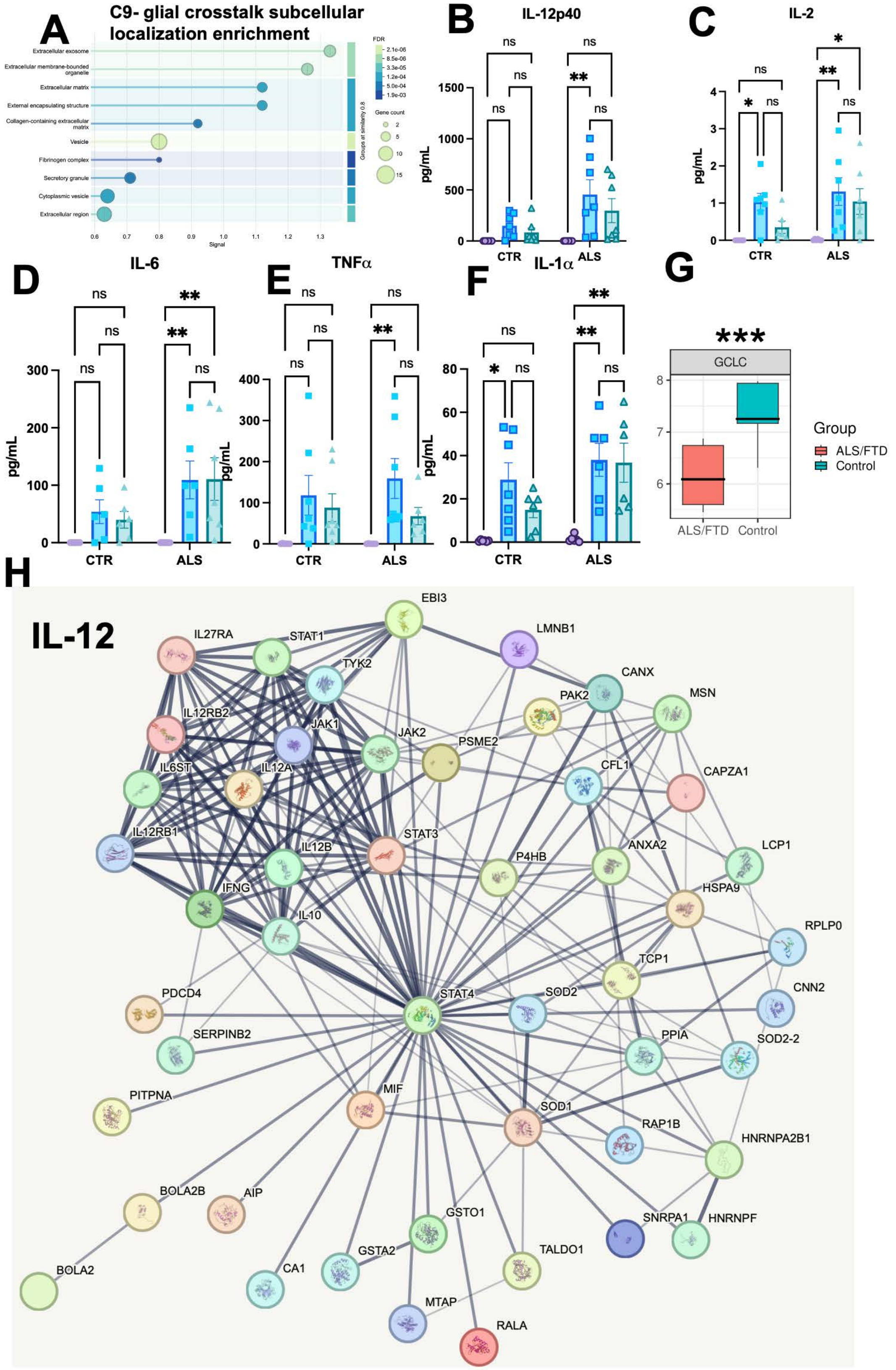


**Figure S3. Inflammatory dysregulation identified in multiple analyses**

A)Subcellular compartment pathway enrichment from CTR vs C9 glial crosstalk.

B-F) Bar graphs of MSD cytokine detection from secreted proteins of stimulated MG. n= 6 CTR and 6 C9 samples. Two-way ANOVA. Protein normalized abundance n=6 CTR and n=6 C9.

G) Boxplot of GCLC expression from bulk-transcriptomics. * $p< 0.05$, ** $p< 0.01$, *** $p<0.001$, **** $p<0.0001$.

H) STRING network projection of IL-12 signaling.

**A**

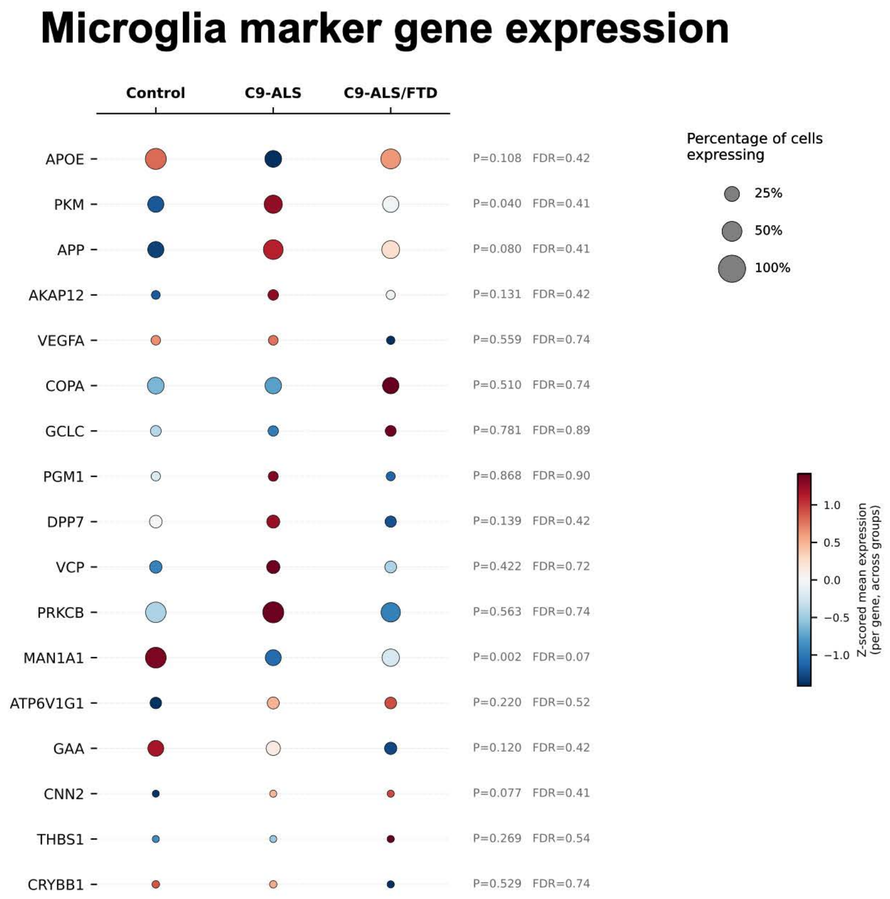


**B**

| Protein/Gene | iPSC C9-MG | C9-ALS postmortem | C9-ALS/FTD postmortem |
|---|---|---|---|
| AKAP12 | = | ↑ | ↑ |
| PKM* | = | ↑ | ↑ |
| VCP | = | ↑ | ↑ |
| ATP6V1G1 | = | ↑ | ↑ |
| CNN2 | ↑ | ↑ | ↑ |
| APP | ↑ | ↑ | ↑ |
| THBS1 | ↑ | ↑ | ↑ |
| APOE | ↓ | ↓ | ↓ |
| CRYBB1 | ↓ | ↓ | ↓ |
| GAA | ↓ | ↓ | ↓ |
| MAN1A1 * | ↓ | ↓ | ↓ |
| VEGFA | ↑ | ↑ | ↓ |
| PGM1 | = | ↑ | ↓ |
| DPP7 | ↓ | ↑ | ↓ |
| GCLC | ↓ | ↓ | ↑ |

*↑ Increased expression/abundance ↓ Decreased expression/abundance = No reported change*

**Figure S4. Microglia expression of hits from snRNA-seq**
A) Expression profiles of hits identified from iPSC glial-crosstalk from snRNA-seq from human frontal cortex microglia.
B) Diagram summarizing changes in iPSC-MG and postmortem C9-nuclei. * p< 0.05 in human tissue.
